# Fetal innate immunity contributes to the induction of atypical behaviors in a mouse model of maternal immune activation

**DOI:** 10.1101/2020.10.09.333815

**Authors:** Eva K. Nichols, Hsiu-Chun Chuang, Matthew T. Davis, Kristina M. Geiger, Rick Z. Li, Madeline L. Arnold, Patrick M. Lin, Rhea Misra, Laurent Coscoy, Kaoru Saijo

## Abstract

Maternal immune activation (MIA) increases likelihood of altered neurodevelopmental outcomes. Maternal cytokines are proposed to affect fetal brain development in mice; however, the contribution of fetal immunity to neurodevelopmental disorders is largely unexplored. Here, we show that MIA mediated by Toll-like receptor 3 (TLR3), but not other TLRs, induces a specific set of behavioral phenotypes including decreased sociability and increased restricted repetitive behavior in offspring. Accordingly, these behavioral phenotypes were absent when offspring were deficient for *Trif*, the downstream adapter molecule of TLR3. Using single-cell RNA sequencing, we identified clusters of border-associated macrophages that were significantly enriched in the fetal brain following TLR3-MIA, and these clusters were diminished in *Trif*^−/−^ fetal brains.Moreover, we found that triggering TLR3-TRIF in offspring can occur through transplacental viral infection, resulting in altered behavioral phenotypes. Collectively, our data indicate that fetal innate immunity contributes to MIA-induced atypical behaviors in mice.

## Introduction

Infections during pregnancy increase the risk of neurodevelopmental disorders (NDDs) in children (Atladottir et al., 2010; Zerbo et al., 2015). To model this epidemiological risk factor in animals, injection of innate immune stimulants, such as toll-like receptor (TLR) ligands, into pregnant wild-type (WT) mice was developed and has been used as the maternal immune activation (MIA) model for NDDs (Smith et al., 2007). MIA results in a variety of behavioral alterations in offspring, including defective sociability (such as decreased social interaction and social memory), increased restricted repetitive behavior, impaired learning and memory, altered levels of anxiety, hyperactivity, and others. Given these observed differences in behaviors, this model is broadly used to study NDDs such as autism spectrum disorders (ASD) (Boksa, 2010; Estes and McAllister, 2016; Patterson, 2011).

To induce MIA, the TLR3 agonist Poly(I:C) or the TLR4 agonist lipopolysaccharide (LPS) are often used in mice, mimicking viral and bacterial infection, respectively (Boksa, 2010; Estes and McAllister, 2016; Patterson, 2011; Solek et al., 2018). TLR3 localizes on the endosomal membrane and signals through its adaptor protein, TRIF, leading to the transcription of anti-viral molecules, such as type I interferons, through the activation of transcription factors IRF3 and IRF7. TLR4 is expressed on the plasma membrane and signals through both the TRIF and MyD88 pathways. MyD88 signaling induces the transcription of pro-inflammatory mediators by activating NF-κB and AP-1 (Supplemental Figure 1A) (Foster and Medzhitov, 2009; Takeda and Akira, 2004). The currently proposed mechanism for MIA-induced impairment of fetal brain development is that TLR ligands activate maternal adaptive immunity. This effect depends on the gut microbiota of dams; particularly, the presence of segmented filamentous bacteria (SFB); which influences the differentiation and activation of Th17 T cells and the production of cytokines such as interleukin (IL)-17A. These maternal cytokines then directly affect fetal brain development (Choi et al., 2016; Kim et al., 2017). So far, it is unknown whether these canonical innate immune TLR signaling pathways, triggered maternally and/or fetally, are required for the developmental outcomes observed in MIA offspring.

Interestingly, a few reports have suggested that MIA alters gene expression in fetal microglia (Ben-Yehuda et al., 2020; Matcovitch-Natan et al., 2016; Mattei et al., 2017). Microglial cells are resident innate immune cells in the brain and spinal cord parenchyma that act as sentinels of infection and injury (Ransohoff and El Khoury, 2015; Saijo and Glass, 2011; Wolf et al., 2017). They originate from yolk-sac primitive macrophages and migrate to the brain at embryonic day 9.5 (E9.5), before the differentiation of neural progenitor cells (NPCs) at E12.5 (Bjornsson et al., 2015; Ginhoux et al., 2010). In addition to surveillance, microglia play essential roles in ensuring normal brain development through the engulfment of dead/dying NPCs, neurons, and synapses to establish proper neural wiring and functions (Brown and Neher, 2014; Paolicelli et al., 2011; Sierra et al., 2010). However, it is unclear whether any specific microglial subsets are affected by MIA, and whether transcriptional changes in microglia contribute to behavior abnormalities.

In addition to microglia, a few subsets of macrophages, such as border-associated macrophages (BAMs), are present in the brain and show distinct patterns of gene expression and localization compared to microglia cells. Like microglia cells, BAMs also originate from yolk-sac primitive macrophages and seed the brain prenatally (Goldmann et al., 2016; Utz et al., 2020). BAMs have been recently proposed to play pathological roles in brain inflammation, contributing to disorders such as experimental autoimmune encephalomyelitis and Alzheimer’s disease (Jordao et al., 2019; Mrdjen et al., 2018). Interestingly, some genes are observed to be expressed in both BAMs and microglia, such as *Fcrls* and *Cx3cr1*, but others are expressed explicitly in only one of these two cell types (Jordao et al., 2019; Mrdjen et al., 2018; Utz et al., 2020; Van Hove et al., 2019). In addition to having unique transcription profiles, microglia and BAMs have distinct patterns of localization. Microglia are localized to the brain parenchyma. BAMs are found in perivascular regions including the meninges and the choroid plexus (CP), where they can be exposed to substances derived from mothers (Mrdjen et al., 2018; Utz et al., 2020; Van Hove et al., 2019). However, BAMs haven’t been previously linked to the development of NDDs. Moreover, we do not know whether observed transcriptional changes are results of fetal microglia or BAMs responding directly to MIA, versus indirect responses to maternal cytokines.

Through genetic and behavioral analyses, we have found that fetal innate immunity triggered by TRIF signaling contributes to the induction of behavior abnormalities such as reduced sociability. Furthermore, using single-cell RNA-sequencing (scRNA-seq), we identified clusters of cells profoundly enriched in wild-type (WT) TLR3-MIA fetal brains that were absent in *Trif*^*−/−*^ TLR3-MIA fetal brains. Using bioinformatics, we identified these cells to be BAMs. Consistent with scRNA-seq data, TLR3-MIA increased the number of BAMs in the CP. We also show that maternal infection with a virus that can cross the placenta can trigger TRIF-dependent behavioral changes in offspring. Based on these results, we propose that transplacental infection may directly trigger fetal TRIF signaling and result in aberrant behaviors. Overall, our data suggests that fetal TRIF signaling in BAMs may play essential roles in the induction of behavioral abnormalities in a mouse model of MIA.

## Results

### Specific TLR ligand-mediated MIA induces discrete sets of behavioral abnormalities in offspring

Both Poly(I:C) (TLR3) and LPS (TLR4)-mediated MIA have been used to induce characteristic behavioral traits, such as decreased sociability, increased restricted repetitive behavior, impaired learning and memory, and other aberrant behaviors in mice (Boksa, 2010; Estes and McAllister, 2016; Patterson, 2011). To compare the effects of these models in our setting, we injected Poly(I:C) and LPS into pregnant wild-type C57BL/6 female mice (WT) at E12.5, a time when NPCs are starting to differentiate (Bjornsson et al., 2015) and ran a panel of assays (Supplemental Figure 1B) to characterize behavior phenotypes in the adult offspring (Figure 1A). Interestingly, we observed reduced sociability, as determined by the 3-chamber social interaction (Figure 1B), reciprocal social interaction, and social transmission of food preference assays (data not shown), as well as increased restricted repetitive behavior using the marble-burying assay (Figure 1C) in Poly(I:C)-MIA offspring, but not in LPS-MIA offspring. We also observed hyperactivity in Poly(I:C)-MIA offspring, but not in LPS-MIA offspring (Supplemental Figure 1C). In contrast, we detected increased anxiety-like behavior in the elevated plus maze assay in LPS-MIA offspring, but not in Poly(I:C)-MIA offspring (Supplemental Figure 1D). Moreover, we observed decreased learning and memory, as determined by the T-maze and fear conditioning assays, in both Poly(I:C)- and LPS-MIA offspring (Supplemental Figure 1E and data not shown). Overall, the behavioral phenotypes induced by MIA differed between Poly(I:C) and LPS treated animals, and are summarized in Figure 1D. Notably, we observed behavioral changes in male offspring of Poly(I:C)-injected dams, but not female littermates (Figure 1E and F).

**Figure 1.**
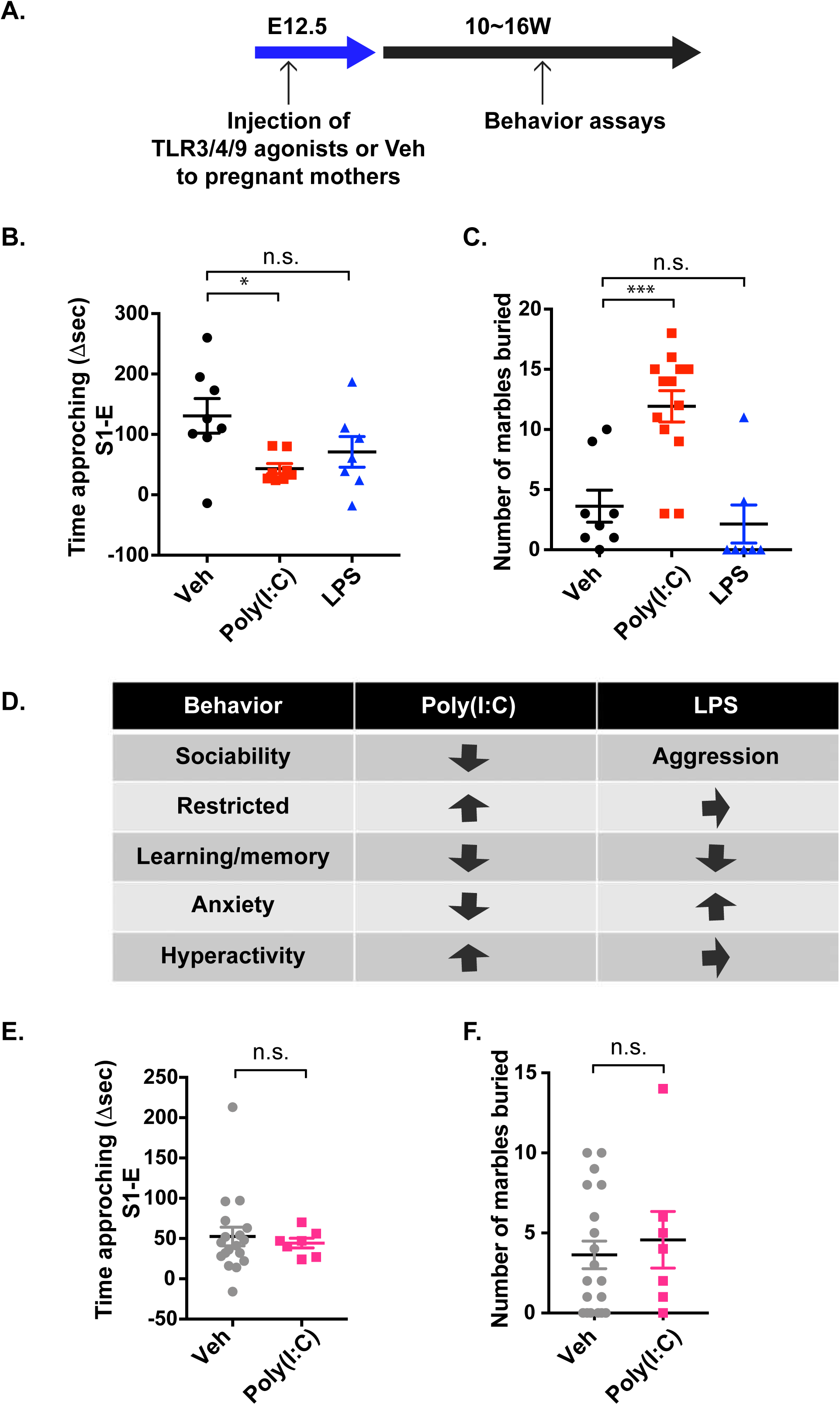
Distinct behavioral phenotypes are induced by Poly(I:C)- and LPS-MIA. **A**. Experimental design is shown. Ligands for TLR3, 4, or 9, or vehicle for control (Veh) were injected into WT pregnant mice at E12.5. Offspring were subjected to behavior assays when they were 10∼16 weeks old. **B**. Social interaction time (Δsec) with a stranger mouse determined by the three-chamber social interaction assay is shown. Male offspring from vehicle control- (Veh, black circles, N=8), Poly(I:C)- (red squares, N=8), and LPS- (blue triangles, N=7) MIA-treated mice were used for this assay. **C**. The degree of restricted repetitive behavior was determined by the number of buried marbles using the marble burying assay. Male offspring from vehicle control (Veh, black circles, N=8), Poly(I:C)- (red squares, N=13), and LPS- (blue triangles N=7) MIA-treated mice were used for this assay. **D**. A summary of behavioral abnormalities induced in male offspring by different MIA models is shown. Arrows indicate behavioral changes compared to vehicle-MIA. **E**. Social interaction time (Δsec) with a stranger mouse determined by the three-chamber social interaction assay is shown. Female offspring from vehicle control- (Veh, silver circles, N=18) and Poly(I:C)- (pink squares, N=7) MIA-treated mice were used for this assay. **F**. The degree of restricted repetitive behavior was determined by the number of marbles buried using the marble burying assay. Female offspring from vehicle control (Veh, silver circles, N=19) and Poly(I:C)- (pink squares, N=7) MIA-treated mice were used for this assay. **B**., **C**., **E**., **F**. Data indicate individual animals and averages. Error bars show S.E.M. * indicates *p*<0.05, *** indicates *p*<0.001, and n.s. indicates no significance.

In addition to the TLR3 and TLR4 ligands, we also tested CpG ODN, a ligand for TLR9, in our MIA model and found that it did not affect sociability (Supplemental Figure 1F) but did decrease restricted repetitive behavior (Supplemental Figure 1G). CpG ODN-MIA also led to impaired learning and memory behaviors in offspring (Supplemental Figure 1H), as well as increased anxiety-like behavior (Supplemental Figure 1I), but not hyperactivity (Supplemental Figure 1J). Interestingly, both LPS- and CpG ODN-MIA offspring exhibited aggressive behavior that we did not observe in Poly(I:C)-MIA offspring (Supplemental Figure 1K). A summary of the behavior abnormalities of CpG ODN-MIA offspring is shown in Supplemental Figure 1L.

Taken together, our data suggest that Poly(I:C)-MIA induces specific behavioral changes in offspring, including reduced sociability and increased restricted repetitive behavior, compared to LPS- and CpG ODN-MIA offspring. Additionally, these behavioral phenotypes are only present in male offspring.

### Fetal TRIF signaling is important for Poly(I:C)-MIA induced behavior changes

Reduced sociability and increased restricted repetitive behaviors are frequently observed in ASD patients and animal models. Previous reports suggested that these aberrant behaviors could be induced by both TLR3 (Poly(I:C))- and TLR4 (LPS)-MIA; however, by our hands, only TLR3-MIA resulted in these behavioral phenotypes. Therefore, next, we decided to investigate the specificity and requirement of maternal TLR3 signaling to induce these behaviors using a genetic approach. TRIF is an essential adaptor molecule for TLR3 signaling (Supplemental Figure 1A) (Uematsu and Akira, 2007; Ullah et al., 2016); therefore, we crossed *Trif* ^−/−^ female with WT male mice, injected the subsequent pregnant mice with Poly(I:C) or vehicle at E12.5, and used the *Trif* ^+/−^ male offspring for our behavioral assays. In this approach, the offspring, but not the mothers, were competent for TRIF signaling (Figure 2A). Restricted repetitive behavior (Figure 2B) and learning and memory (Supplemental Figure 2A) did not differ between offspring from Poly(I:C) or vehicle injected *Trif* ^−/−^ dams. However, surprisingly, we still observed decreased sociability (Figure 2C), lower anxiety-like behavior (Supplemental Figure 2B), and hyperactivity (Supplemental Figure 2C) in these mice.

**Figure 2.**
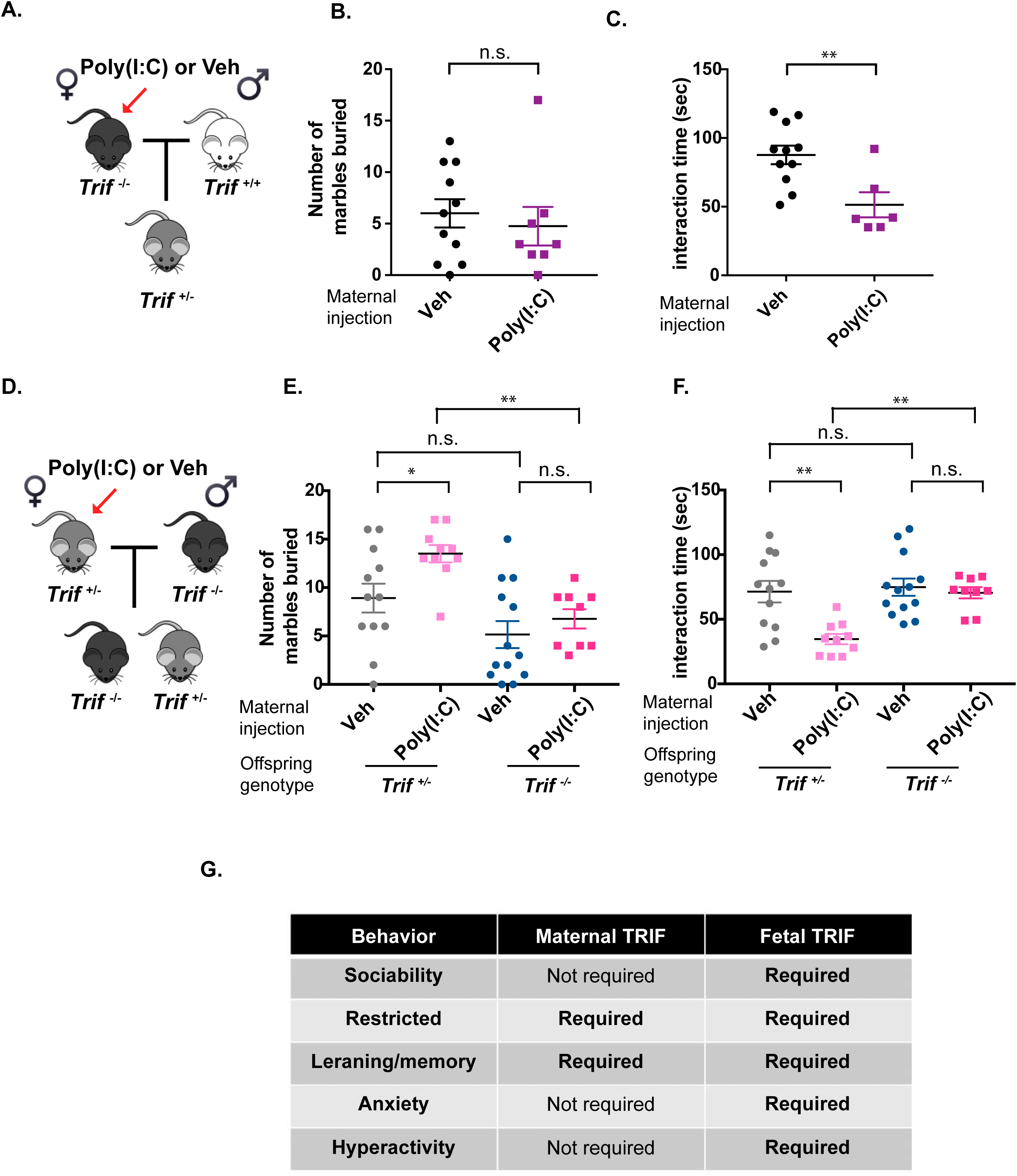
MIA offspring exhibit maternal and fetal *Trif*-dependent behavioral phenotypes. **A**. Experimental design is shown. *Trif*^−/−^ female mice were crossed with WT male mice to generate *Trif*^*+/−*^ offspring. Poly(I:C) or vehicle (Veh) was injected into pregnant mice at E12.5 and behavior assays were performed on offspring at 10-16 weeks of age, as shown in Figure 1A. **B**. The degree of restricted repetitive behavior was determined by the number of marbles buried using the marble burying assay. Offspring from vehicle control- (Veh, black circles, N=11) and Poly(I:C)- (purple squares, N=8) MIA-treated mice were used for this assay. **C**. Social interaction time (sec) with a stranger mouse determined by the reciprocal social interaction assay is shown. Offspring from vehicle control- (Veh, black circles, N=11) and Poly(I:C)- (purple squares, N=6) MIA-treated mice were used for this assay. **D**. Experimental design is shown. *Trif*^*+/−*^ female mice were crossed with *Trif*^−/−^ male mice, resulting in both *Trif*^−/−^ and *Trif*^+/−^ offspring. Poly(I:C) or vehicle (Veh) was injected into pregnant mice at E12.5 and behavior assays were performed when offspring reached 10-weeks of age, as shown in Figure 1A. **E**. The degree of restricted repetitive behavior was determined by the number of marbles buried using the marble burying assay. *Trif*^+/−^ offspring from vehicle control- (Veh, silver circles, N=12) and Poly(I:C)- (pink squares, N=10) MIA-treated mice, as well as *Trif*^−/−^ offspring from vehicle control- (Veh, blue circles, N=13) and Poly(I:C)- (strawberry squares, N=9) MIA-treated mice were used for this assay. **F**. Social interaction time (sec) with a stranger mouse determined by the reciprocal social interaction assay is shown. *Trif*^+/−^ offspring from vehicle control- (Veh, silver circles, N=12) and Poly(I:C)- (pink squares, N=10) MIA-treated mice, as well as *Trif*^−/−^ offspring from vehicle control- (Veh, blue circles, N=13) and Poly(I:C)- (strawberry squares, N=9) MIA-treated mice were used for this assay. **B**., **C**., **E**., **F**. Data indicate individual animals and averages. Error bars show S.E.M. * indicates *p*<0.05, ** indicates *p*<0.01, and n.s. indicates not significant. **G**. A summary of the maternal and fetal TRIF signaling requirements for the behavior abnormalities indicated is shown.

We thought of two possibilities to explain why Poly(I:C)-MIA partially induced altered behaviors in the offspring of Poly(I:C)-injected *Trif* ^−/−^ dams. The first possibility is that another sensor, such as the RIG-I/MAVS pathway (Seth et al., 2005; Yoneyama et al., 2004), is required for sensing Poly(I:C) in dams. The second possibility is that the fetal innate immune system may contribute to the induction of altered behaviors. To test the latter possibility, we bred *Trif* ^+/−^ females with *Trif* ^−/−^ males, injected Poly(I:C) into the pregnant mice at E12.5, and used the *Trif* ^−/−^ and *Trif* ^+/−^ offspring for our behavior assays. In this approach, the mother was competent for TRIF signaling while the homozygous null offspring were not (Figure 2D). Poly(I:C)-MIA increased restricted repetitive behavior and decreased sociability in the *Trif* ^+/−^ offspring of *Trif* ^+/−^ dams (Figure 2E and F). However, Poly(I:C)-MIA failed to affect behaviors in offspring deficient for *Trif* (Figure 2E and F). Likewise, we did not observe impaired learning and memory (Supplemental Figure 2D) or lower anxiety levels (Supplemental Figure 2E) in *Trif* ^−/−^ offspring. We did not observe any differential behaviors between *Trif* ^+/−^ and *Trif* ^−/−^ offspring when their mothers were injected with the vehicle control (Figure 2E and F). Interestingly, Poly(I:C)-MIA not only did not induce hyperactivity in *Trif* ^−/−^ offspring, but *Trif* ^−/−^ offspring of treated dams were less active than vehicle controls (Supplemental Figure 1F).

Overall, our data suggests that both maternal and fetal TRIF-signaling contribute to behavioral phenotypes induced by TLR3/Poly(I:C)-MIA, as summarized in Figure 2G. Importantly, fetal TRIF-signaling is required for the induction of multiple behavior abnormalities, including reduced sociability, restricted repetitive behavior, hyperactivity, and impaired learning and memory.

### Characterization of fetal brain myeloid cells by single-cell RNA sequencing following MIA

Our behavioral data indicated that fetal innate immunity is required for the induction of many altered behaviors in the offspring of Poly(I:C)-injected mothers, implying that fetal immune cells were directly responding to the TLR ligand (Figure 2E-G). To identify the fetal responder cells, we used single-cell RNA sequencing (scRNA-seq). For these studies, we injected Poly(I:C) or vehicle control into pregnant WT or *Trif* ^−/−^ mice at E12.5, and fetal brains were harvested 24 hours later, including two separate litters for each combination of treatment and genotype. CD45^+^CD11b^+^ myeloid cells were sorted from E13.5 fetal brains, and loaded onto the 10X Genomics scRNA-seq platform (Figure 3A).

**Figure 3.**
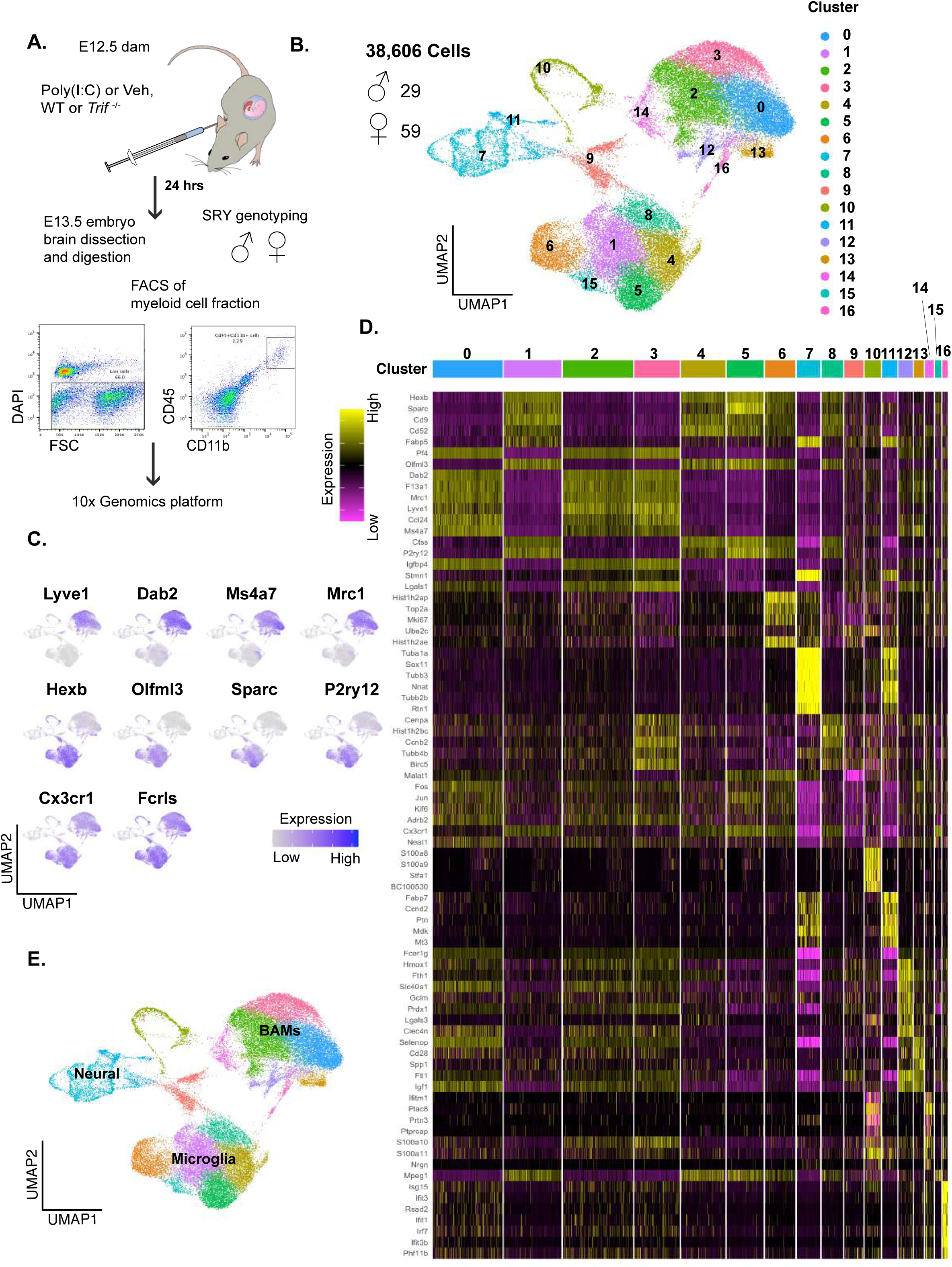
Identification of clusters by scRNA-seq. **A**. Experimental design is shown. WT female were crossed to WT male and *Trif*^−/−^ female were crossed to mice *Trif*^−/−^ male mice. Pregnant mice were injected with Poly(I:C) or vehicle (Veh) at E12.5 and fetal brains were harvested 24 hours later. Both male and female brains were dissected, dissociated, and cells were sorted to isolate the CD11b^+^CD45^+^ population. Libraries were generated and run on the 10X Genomics platform. **B**. UMAP plot of all the cells from two genotypes (WT and *Trif*^−/−^ mice) and two treatments (Poly(I:C) or Veh) is shown. A total of 17 clusters were identified. **C**. UMAP plot of the canonical markers for BAMs (top row), microglia (middle row), and both BAMs and microglia (bottom row). **D**. Heatmap of genes with the highest absolute value log fold change returned from differential expression analysis between each cluster and the rest of the cells in the dataset. Each row is a gene, and each column in each row belongs to a cell within a cluster. Names of the marker genes are on the left. The Cluster IDs are color coded at the top of the heatmap. **E**. Identities of Clusters shown in Figure 3B.

The resulting 8 batches are summarized in Supplemental Figure 3A. We recovered a total of 49,865 cells from 29 male and 59 female fetal brains, and after running the doublet finder algorithm (McGinnis et al., 2019), we were left with 38,606 cells. With these cells, we achieved a median ∼60,000 reads per cell, an average median capture of 11,080 UMIs (unique molecular identifiers), and ∼2000 unique genes per cell per batch with less than 15% mitochondrial gene content per cell per batch (Supplemental Figure 3B). With principal component analysis (PCA) and k-means clustering, we obtained 17 clusters based on the dimensionally reduced gene expression data of these cells (Figure 3B) (details in the methods). In our analysis, we found that 35,191 cells were positive for *Fcrls*, a marker gene for both border-associated macrophages (BAMs) and microglial cells (Figure 3C and Supplemental Figure 3C). BAMs have been found to highly express genes such as *Lyve1*, Dab2, *Ms4a7*, and *Mrc1*, while microglia highly express genes including *Hexb, Olfml, Sparc, and P2ry12* (Jordao et al., 2019; Mrdjen et al., 2018; Utz et al., 2020; Van Hove et al., 2019). Consistent with previous literature, we detected these BAM and microglial gene signatures in the *Fcrls*^*+*^ cell fractions (Figure 3C and Supplemental Figure 3D and E). Based on the expression of *Fcrls* and these canonical marker genes, we defined Clusters 0, 2, 3, 12, 13, 14, and 16 as BAMs, and Clusters 1, 4, 5, 6, 8, and 15 as microglia (Figure 3C-E and Supplemental Figure 3D and E). Clusters 9 and 10 were positive for *Fcrls* expression and exhibited both BAM and microglial-like signatures; thus, their identities could not be determined from marker gene identification alone (Supplemental Figure 3C-E). We also found *Fcrls*^−^ cells (Clusters 7 and 11), which were likely neural cells (Figure 3D and E).

### Identification of putative fetal innate immune responder cells in the fetal brain

We found that *Fcrls*^*+*^ cells in Clusters 9 and 10 were significantly enriched in Poly(I:C)-MIA fetal brains (Figure 4A and B and Supplemental Figure 4A). We also found that these clusters were primarily composed of WT cells, with a minor contribution from *Trif* ^−/−^ cells (Figure 4C). Given that Clusters 9 and 10 were largely derived from one batch (Supplemental Figure 4B), we sought to validate them further. To determine whether the separation of Clusters 9 and 10 was due to technical factors that contributed to batch effects, such as differential UMIs captured, we downsampled every cell to 2500 random UMIs (Supplemental Figure 4C). Both clusters remained after downsampling, suggesting that they are biologically relevant (Dixit et al., 2016; Luecken and Theis, 2019; Vieth et al., 2019). Interestingly, we did not observe any significant differences between the number of male and female cells in Cluster 10 and others; however, Cluster 9 was predominantly comprised of male cells (Supplemental Figure 4D and E). These data suggest that the cells in Clusters 9 and 10 may be the putative cells responsible for fetal TLR3-TRIF signaling in response to MIA.

**Figure 4.**
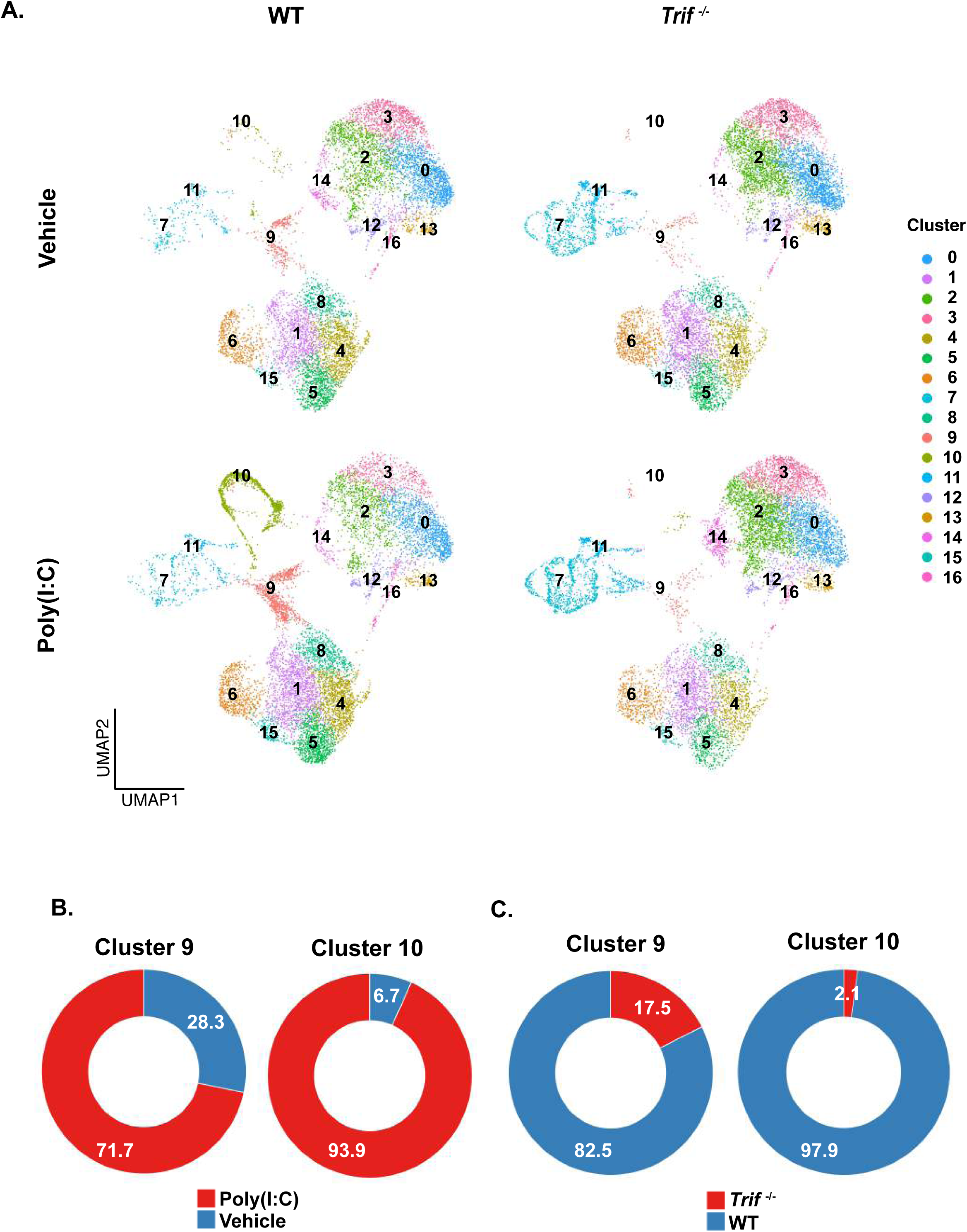
Clusters 9 and 10 are enriched in fetal brain cells from the offspring of WT Poly(I:C)-MIA-treated mice. **A**. UMAP plots of fetal brains isolated from WT (left) and *Trif*^−/−^ (right) offspring of vehicle (top) and Poly(I:C)- (bottom) MIA-treated mice are shown. The Clusters are color-coded and shown on the right. **B**. Pie charts representing the contributions of vehicle- and Poly(I:C)-MIA treated cells to Clusters 9 (left) and 10 (right) are shown. **C**. Pie charts depicting the contributions of *Trif*^−/−^ and WT cells to Clusters 9 (left) and 10 (right) are shown.

### Fetal myeloid responder cells to Poly(I:C)-induced MIA are likely BAMs

UMAP plots indicated that Clusters 9 and 10 were clearly segregated from major clusters of BAMs and microglia (Figure 3B). Since these clusters expressed both BAM and microglial marker genes (Supplemental Figure 3D and E), we performed a few more analyses to determine their precise identities. First, we used an RNA velocity (Velocyto) approach to predict how cells in these clusters are progressing towards future transcriptional states in pseudotime, and determine whether trajectories toward future patterns of gene expression lead to BAM- or microglia-like cell states (La Manno et al., 2018). We found that the BAM and microglial sub-clusters trended toward a steady cell state represented by clusters 0 and 1, respectively (Figure 5A). Clusters 0 and 1 were G1 cells, while other sub-clusters were undergoing active mitosis in G2/M and S stages (Supplemental Figure 5A). These results suggest that other sub-clusters may be actively proliferating cells filling the BAM and microglial niches. From here on, we refer to Cluster 0 as “steady state BAMs” and Cluster 1 as “steady state microglia”. While our random control did not show any clear pattern (Supplemental Figure 5B), Cluster 10 trended toward a BAM-like cell state (Figure 5A). Some Cluster 9 cells seemed to be at a steady state, while other cluster 9 cells also moved toward BAMs; these data indicate that, irrespective of origin, Poly(I:C)-MIA responder cells that are Clusters 9 and 10 are likely to be BAMs.

**Figure 5.**
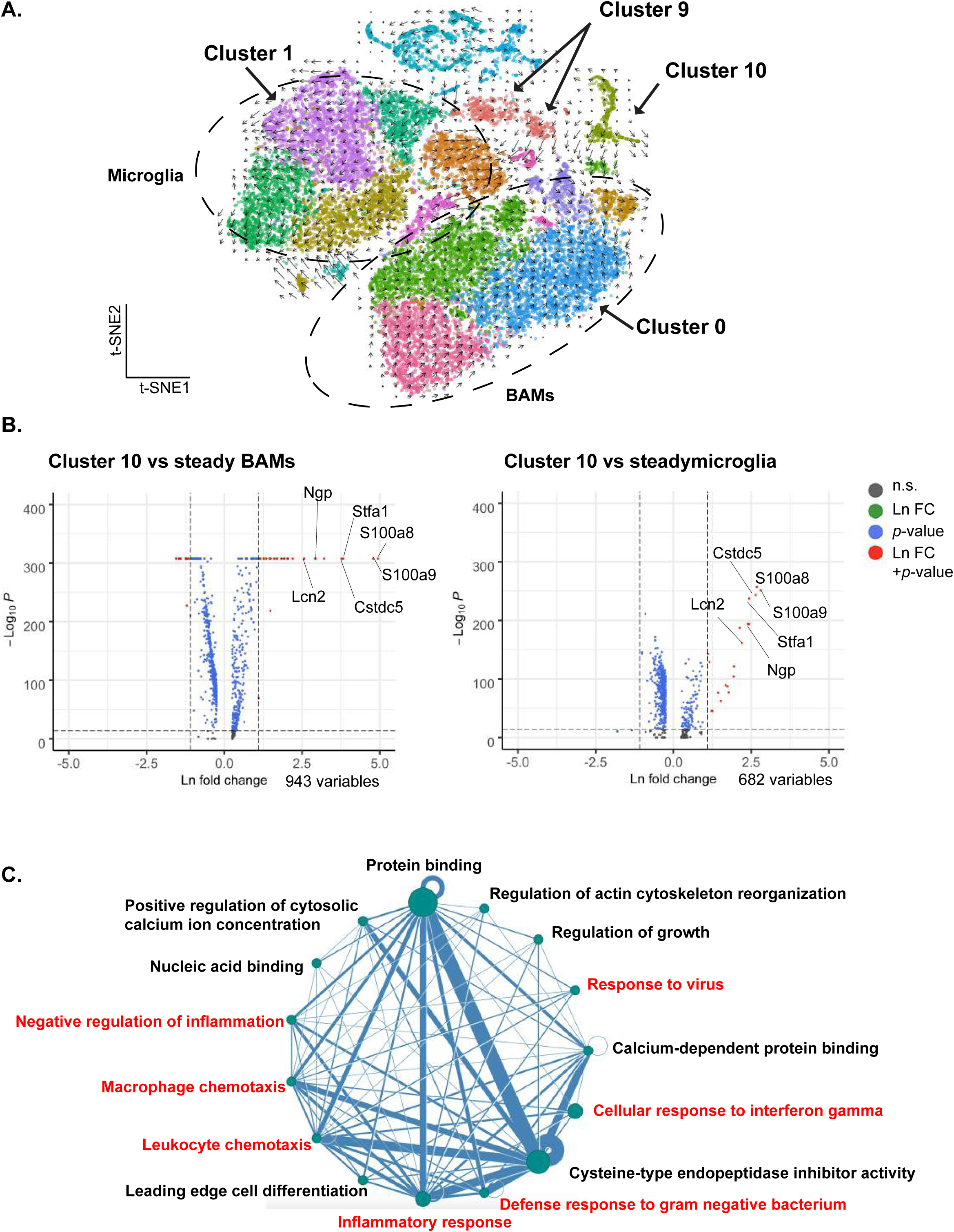
Cluster 10 cells trend toward a BAM-like cell state and are enriched in genes regulating inflammation. **A**. RNA velocity analysis performed using the Velocyto program was dimensionally reduced and is presented as a t-SNE plot. Dashed circles indicate the microglial and BAM cell neighborhoods, as defined previously. The direction of the arrows represents the predicted movement of cells in pseudotime based on the ratios of spliced and unspliced RNAs in each cell. Cluster 1 represents a steady cell state within the microglial cell neighborhood, while Cluster 0 represents that within the BAMs. Only the WT portion of the dataset is presented here. **B**. Volcano plots representing the differential gene expression observed in Cluster 10 compared to Clusters 0 (left) and 1 (right). Orange dots represent differentially expressed genes in Cluster 10 compared to Clusters 0 and 1 (upregulated gene cutoff is log2fold change > 1.5 and Bonferonni adjusted p value (p adj) < 0.01; downregulated gene cutoff is log2fold change < −1.5 and *p adj* < 0.05). **C**. Network of biological pathways inferred by patterns of co-expression amongst the most highly expressed markers of cluster 10. Each node represents a set of >= 2 genes with shared gene ontology. The width of each connection is determined by the Pearson’s r value of correlated expression between each pair of genes across all cells in the cluster. For two nodes, the average r value of each pair of genes between those nodes determines the width of the connection. Inflammation related categories are highlighted in red.

Next, we determined the gene expression patterns characterizing Clusters 9 and 10. With differential expression analysis, we identified genes upregulated in Cluster 9 and Cluster 10 compared to clusters of steady state BAMs and microglia (Figure 5B). Of note, we found high expression of *S100a8* and *S100a9* in Cluster 10 (Figure 5B), which have been previously shown to be upregulated in activated BAMs and microglia (Jordao et al., 2019; Xia et al., 2017). Using the network analysis program MTGO-sc (Nazzicari et al., 2019), we confirmed that highly coexpressed markers of Cluster 10 form pathways associated with inflammatory response (Figure 5C). Specifically, those pathways included defense response to bacteria, chemotaxis, neutrophil aggregation, and protein binding (including complements and chemokine receptors; Figure 5C and Supplemental Figure 5C) (Nazzicari et al., 2019). While cells in Cluster 10 had upregulation of genes related to inflammation, cells in Cluster 9 were characterized by downregulation of genes canonically expressed in brain myeloid cells compared to steady BAMs and microglia (Figure 3D and Supplemental Figure 5D). Because the network analysis relies on strong co-expression of positive markers it was not feasible to perform on Cluster 9.

Because Cluster 10 showed strong positive biomarker expression, we sought to validate Cluster 10 *in situ* as one population of fetal BAMs which respond to Poly(I:C)- MIA in a *Trif*-dependent manner.

### Fetal CD206^+^ cell numbers are increased in the choroid plexus with Poly(I:C)-MIA

To further investigate whether cells in Cluster 10 that are the putative Polyl(I:C)-MIA responder cells identified in the scRNA-seq analysis, we used immunofluorescence to find the cells expressing Cluster 10 marker genes. For these studies, we used CD206 and P2Y12 as markers for BAMs and microglia, respectively, and we chose *S100a8* and *S100a9* because they were the most highly expressed marker gene defining Cluster 10 (Figure 5B and Figure 6A). Our results showed that the numbers of CD206^+^ cells were significantly increased in the CP of fetal brains (Figure 6B) following Poly(I:C)-MIA. These CD206^+^ cells were also found to be positive for S100a8/9 (Figure 6B and C), with the fluorescence intensity of S100a8/9 increasing with MIA (Figure 6D). Since it has been reported that BAMs primarily localize to the CP and meninges, we also looked for BAMs in the meninges of the fetal brain. Based on our imaging, we did not observe any increase in numbers of CD206^+^ cells in the meninges with Poly(I:C)-MIA (Supplemental Figure 6A and B).

**Figure 6.**
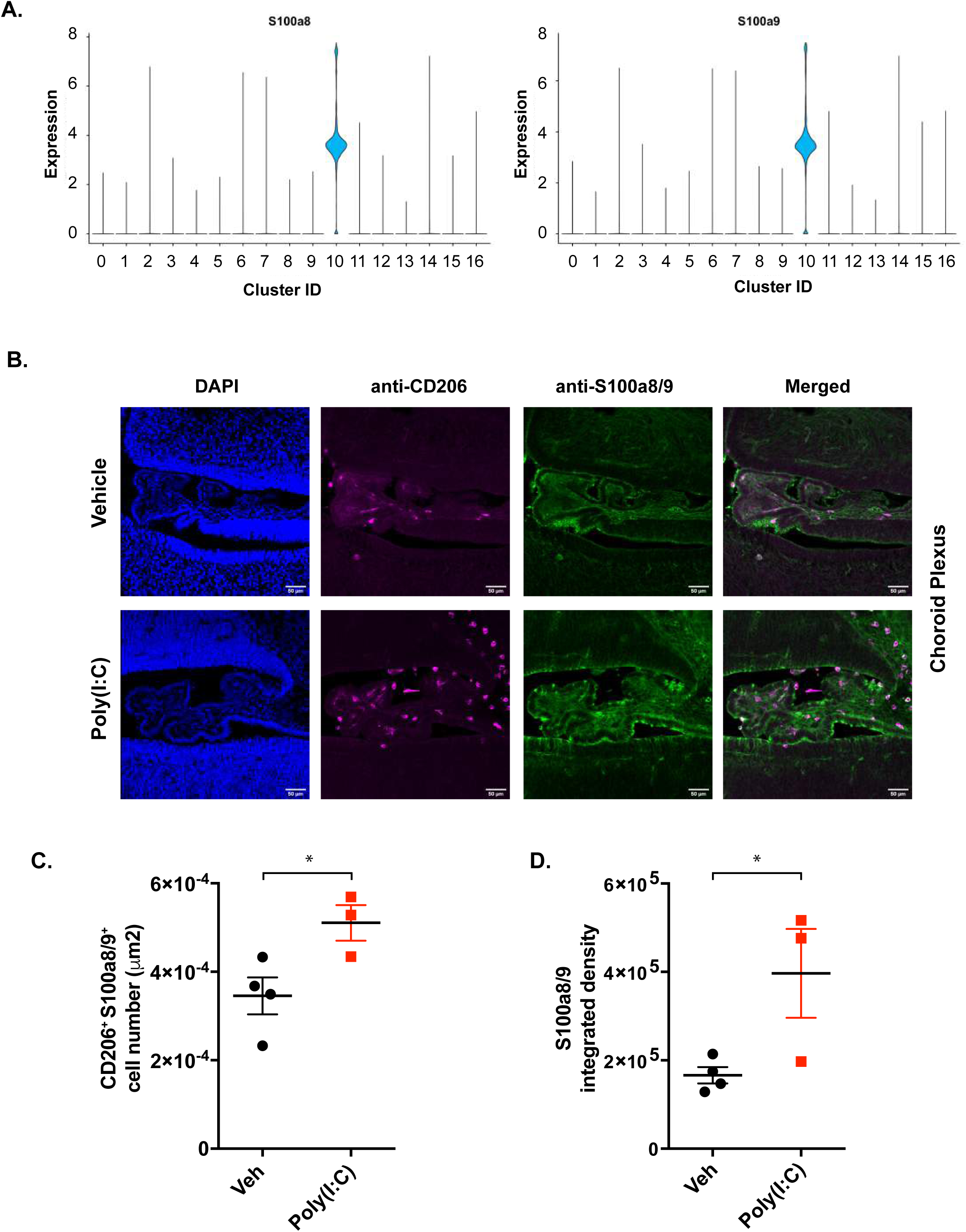
Increased numbers of CD206^+^S100a8/9^+^ cells in the choroid plexus. **A**. Violin plot of S100a8 (left) and S100a9 (right) expression is shown. The width of each violin represents the number of cells at any given expression value. **B**. Representative imaging of fetal brain slices from offspring 24 hours after Poly(I:C)- (bottom panel) or vehicle- (top panel) MIA were stained with anti-CD206, and anti-S100a8/9 antibodies in the CP. Error bars indicate 50 µm. **C**. Quantitation of the number of CD206^+^S100a8/9^+^ cells in the CP from B, Veh (black circle, N=4) and Poly(I:C)-MIA (red squire N=3) is shown. **D**. Quantitation of integrated density of S100a8/9+ signal in the CD206+ cells of CP from B is shown. **C and D**. Each dot corresponds to an individual fetal mouse, and indicates the average cell number or integrated density in 4 to 10 serial sections of the CP and error bars indicate S.E.M. Asterisk (*) indicate that p-value<0.05.

Although microglia are not predominantly localized to the CP, we did stain fetal brains with an anti-P2Y12 antibody to identify microglia in this region. We did not observe increased numbers of fetal microglia in the CP following Poly(I:C)-MIA (Supplemental Figure 6C and D). Moreover, since P2Y12^+^ microglial cells can be found in the brain parenchyma (Mrdjen et al., 2018; Utz et al., 2020; Van Hove et al., 2019), we also counted the numbers of microglia in the fetal cortex of control and Poly(I:C)-MIA offspring. We found no significant changes in microglial cell numbers in the fetal cortex following MIA (Supplemental Figure 6E and F). Overall, our imaging data suggests that the numbers of S100a8/9^+^CD206^+^ BAMs in the CP region are significantly increased in the brains of offspring from Poly(I:C)-treated dams, which is consistent with our scRNA-seq analysis.

### MHV-68 passes through the placenta and induces behavior abnormalities in offspring

Finally, we asked how fetal TLR3-TRIF signaling could be activated in the brain of offspring. We hypothesized that the fetal innate immune system could be activated directly by transplacental infection, and that this route of infection is necessary to produce the behavioral phenotypes associated with fetal innate immune activation. To test this hypothesis, we chose to infect pregnant mice with murine cytomegalovirus (MCMV) and murine gammaherpesvirus-68 (MHV-68, related to human Epstein-Barr virus) as models. Both MCMV and MHV-68 trigger TLR3 upon infection of mice, in spite of being double-stranded DNA viruses (Bussey et al., 2014; Cardenas et al., 2010; Krmpotic et al., 2003; Shen et al., 2018; Simas and Efstathiou, 1998). It is also well established that MCMV does not pass through the placenta in mice (Cardenas et al., 2010; Johnson, 1969); however, MHV-68 may pass through the placenta, though this has been debated (Cardenas et al., 2010; Johnson, 1969; Stiglincova et al., 2011). Therefore, to determine whether MHV-68 passes through the placenta, we infected pregnant WT mice with MHV-68 at E12.5 and tested whether viral DNA was detected in fetal organs 5 days after maternal injection. Our results showed that MHV-68 viral DNA was present in fetal organs, including the brain (Supplemental Figure 7A). Having validated our approach, we used MCMV and MHV-68 to test our hypothesis. Following injection of these viruses into pregnant WT females at E12.5 (Figure 7A), we observed decreased sociability (Figure 7B and C), and increased restricted repetitive behavior (Figure 7D and E) in the offspring of MHV-68-injected dams, but not in the offspring of MCMV-injected dams, suggesting that the fetal innate immune system is indeed activated by transplacental infection, and that this infection is necessary to induce behavioral changes in offspring.

**Figure 7.**
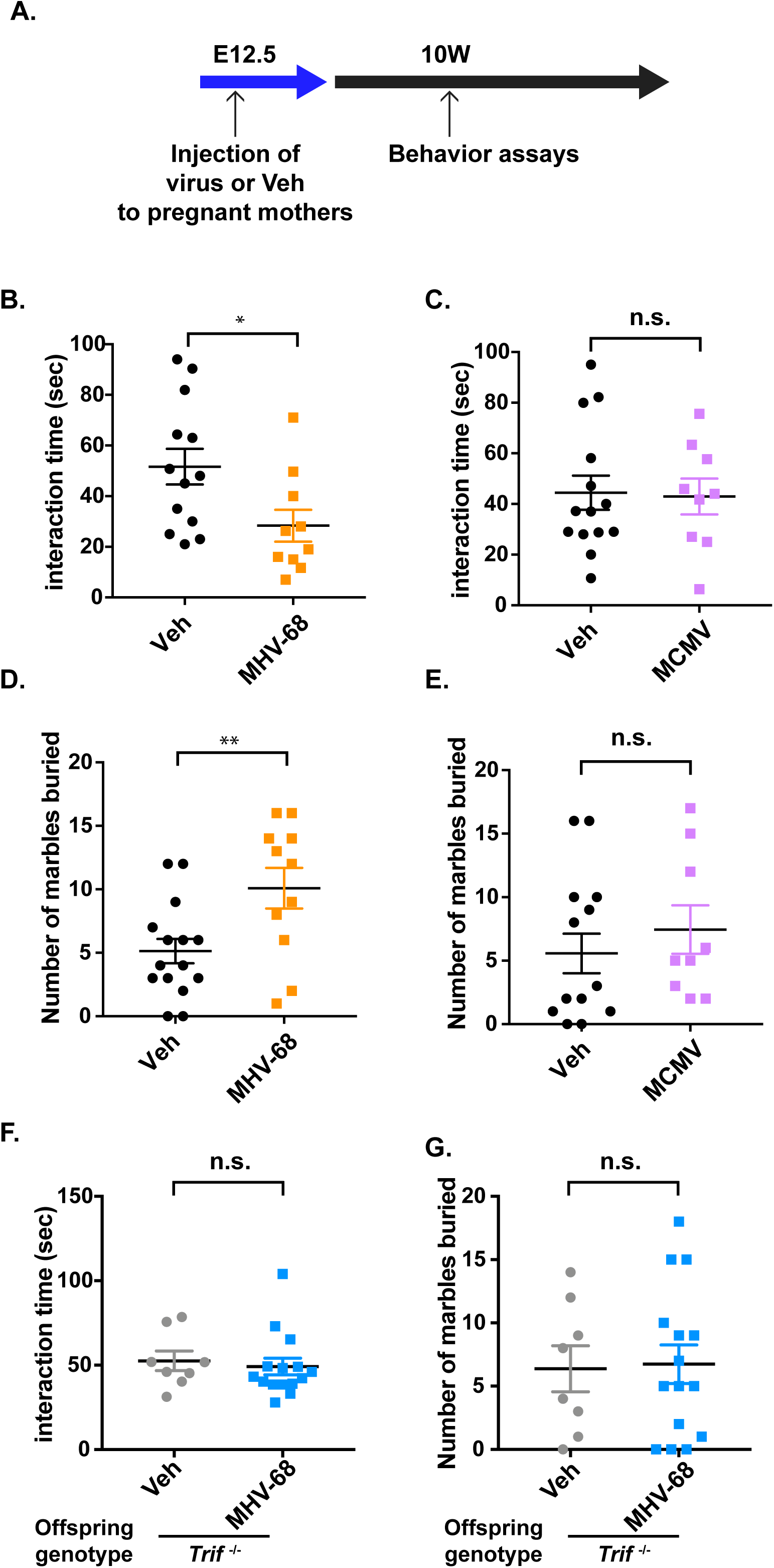
Transplacental infection induces behavior abnormalities. **A**. Experimental design is shown. WT or *Trif*^−/−^ female mice were injected with virus at E12.5 and their offspring were used for behavior assays at 10-16 weeks of age. **B**. Social interaction time (sec) with a stranger mouse determined by the reciprocal social interaction assay is shown. Offspring from vehicle control- (Veh, black circles, N=13) and MHV-68-infected mothers (orange squares, N=10) were used for this assay. **C**. Social interaction time (sec) with a stranger mouse determined by the reciprocal social interaction assay is shown. Offspring from vehicle control- (Veh, black circles, N=14) and MCMV-infected mothers (purple squares, N=9) were used for this assay. **D**. The degree of restricted repetitive behavior was determined by the number of marbles buried using the marble burying assay. Offspring from vehicle control- (Veh, black circles, N=15) and MHV-68-infected mothers (orange squares, N=11) were used for this assay. **E**. The degree of restricted repetitive behavior was determined by the number of marbles buried using the marble burying assay. Offspring from vehicle control- (Veh, black circles, N=14) and MCMV-infected mothers (purple squares, N=9) were used for this assay. **F**. Social interaction time (sec) with a stranger mouse determined by the reciprocal social interaction assay is shown. *Trif*^−/−^ offspring from vehicle control- (Veh, gray circles, N=8) and MHV-68-infected mothers (blue squares, N=15) were used for this assay. **G**. The degree of restricted repetitive behavior was determined by the number of marbles buried using the marble burying assay. *Trif*^−/−^ offspring from vehicle control- (Veh, gray circles, N=8) and MHV-68-infected mothers (blue square, N=15) were used for this assay. **B**.**-G**. Data indicate individual animals and averages. Error bars show S.E.M. * indicates *p*<0.05, ** indicates *p*<0.01, and n.s. indicates not significant.

Based on the above results, we hypothesized that transplacental infection triggers fetal TLR3-TRIF signaling to induce aberrant behaviors in offspring. To test this hypothesis, we infected pregnant *Trif* ^−/−^ female crossed to *Trif* ^−/−^ male mice with MHV-68 and examined behavior of their *Trif* ^−/−^ offspring (Figure 7A). Since reduced sociability is dependent upon fetal TLR3-TRIF signaling (Figure 2F) and increased restricted repetitive behavior is dependent upon both fetal and maternal TLR3-TRIF signaling (Figure 2G), we expected that MHV-68-induced MIA would not lead to behavior abnormalities in *Trif* ^−/−^ offspring. Indeed, we confirmed that these behavioral abnormalities were not induced in *Trif* ^−/−^ offspring following MHV-68-MIA (Figure 7F and G).

Overall, our data suggests that the behavioral phenotypes observed in offspring following Poly(I:C)-MIA may be induced as a result of viral sensing by fetal BAMs. Our working model is shown in Supplemental Figure 7B.

## Discussion

The etiologies of NDDs are complex because both genetic and environmental factors are involved. Mouse models of MIA have been used to study the contribution of environmental factors, demonstrating the importance of maternal Th17 T cells in the establishment of NDDs (Choi et al., 2016; Kim et al., 2017). Since the activation of innate immunity by various pathogens can be an upstream event for adaptive immunity (Iwasaki and Medzhitov, 2015), we investigated the contribution of MIA to associated behavioral phenotypes in offspring using different TLR ligands. Our results showed that signal-dependent MIA can induce discrete behavioral phenotypes. More specifically, Poly(I:C)-MIA triggers a TRIF-dependent decrease in sociability and increase in restricted repetitive behavior, while LPS- and CpG ODN-MIA cause increased aggression and anxiety levels.

LPS, a ligand for TLR4, utilizes the signaling adaptors TRIF and MyD88, while CpG ODN recognition by TLR9 triggers only the signaling cascade mediated by MyD88. Based on previous findings in the literature (Solek et al., 2018), we expected LPS-MIA to induce behavioral phenotypes similar to Poly(I:C)-MIA (Boksa, 2010; Patterson, 2011), but, surprisingly, LPS induction was more similar to CpG ODN-MIA. Our genetic approach using *Trif* ^*−/−*^ dams confirmed that restricted repetitive behavior and defective learning and memory are dependent upon the maternal TLR3-TRIF pathway. However, strikingly, we found that fetal TRIF signaling is also essential for the induction of behavior phenotypes, including decreased sociability, restricted repetitive behavior, impaired learning and memory, lower anxiety-like behavior, and hyperactivity (Figure 2E-F, Supplemental Figure 2D-F). To our knowledge, this is the first work to identify a distinct contribution of fetal immunity to the specific behavioral alterations observed in MIA offspring.

To better understand how fetal immunity is activated in a TLR3-TRIF-mediated mouse model of MIA, we investigated viral infection as a possible trigger for NDD via the fetal TLR3-TRIF pathway. For these studies, we opted to use viruses that are known to trigger TLR3 and that do or do not cross the placenta. Unlike human CMV, which crosses the placenta and causes congenital anomalies (Britt, 2018), it has been well documented that MCMV does not cross the placenta but triggers TLR3 (Cardenas et al., 2010; Johnson, 1969). MHV-68 has been published as a virus that both can (Stiglincova et al., 2011) and cannot (Cardenas et al., 2010) cross the placenta, but in our system, we observed that the virus can indeed cross the placenta and infect the fetus (Supplemental Figure 7A). Although MHV-68 is a DNA virus and TLR9 plays a significant role in clearing the infection and avoiding viral latency, it is also known to trigger TLR3 at the initial infection (Bussey et al., 2014; Shen et al., 2018). Our results showed that MHV-68, but not MCMV, induced a *Trif*-dependent reduction in sociability and an increase in restricted repetitive behavior, suggesting that fetal TLR3-TLR activation through transplacental infection, in addition to maternal cytokines, contribute to the behavioral phenotypes observed in the Poly(I:C)-MIA model.

To identify the fetal innate immune cells that are responsible for TRIF-signaling following MIA, we performed scRNA-seq and found that the fetal responder cells (Clusters 9 and 10) were prominent in Poly(I:C)-MIA fetal brain among *Fcrls*^*+*^ cells (Figure 4A). Furthermore, the number of cells in these clusters markedly increased in WT, but not in *Trif*^*−/−*^, fetal brains. Interestingly, cells in Cluster 9 are primarily characterized by downregulation of marker genes associated with BAMs and microglia, such as (*Cx3cr1* and *Mrc1*) (Jordao et al., 2019; Mrdjen et al., 2018; Utz et al., 2020; Van Hove et al., 2019), AP-1 transcriptional family (*Fos, Jun*, and *JunB*) (Kaminska et al., 2016), chemokines (*Ccl2, Ccl3, Ccl4*, and *Ccl7*) (Mantovani et al., 2004) and regulators of macrophage polarization and activation (Klf6 and Malat1) (Cui et al., 2019; Date et al., 2014). The transcriptional changes observed in Cluster 9 do not correlate with defined sub-macrophage cells such as M1 and M2 polarized macrophages (Lawrence and Natoli, 2011) or myeloid-derived suppressor cells (Alshetaiwi et al., 2020). Because we analyzed the RNA expression 24 hours after Poly(I:C) stimulation (Park et al., 2006), we may have detected transient cell populations involved in the induction or resolution of inflammation. Given that we observed behavioral phenotypes only in male offspring (Figure 1), it is noteworthy that cells in Cluster 9 derive largely from male fetuses (Supplemental Figure 4E). The differential regulation of inflammation between male and female may contribute to the male bias of behavioral alterations in MIA model. Analysis of different kinetics may help to determine the identity of cells in Cluster 9 and potentially sex dimorphism in the model.

In contrast, pathway analysis of the genes enriched in Cluster 10 showed that these genes regulate inflammation (Figure 5B and C, Supplemental Figure 5C). Although cells in Cluster 10 expressed both BAM and microglial marker gene (Supplemental Figure 3D and E), Velocyto analysis indicated that the cells in Cluster 10 are undergoing transcriptional changes which would likely resolved toward a BAM-like cell state in the G1 stage of the cell cycle, rather than a microglial cell state (Figure 5A and Supplemental Figure 5A). Our analysis indicated that these cells have upregulation of biological pathways involved in chemotaxis (Figure 5C), raising the possibility that BAMs activated by Poly(I:C) migrate to the CP by chemotaxis. Further studies are needed to test whether the increase in BAMs we observed in the CP is due to altered patterns of migration. Since our time point of scRNA-seq analysis was 24 hours after Poly(I:C) induction, which is relatively late and is past the peak of pro-inflammatory gene expression, we speculate that we would see a more consistent representation of fetal responder cells if we harvested the fetal brain cells at an earlier time point.

A previous report indicated that BAMs might contribute to neuroinflammatory diseases in adult, such as Alzheimer’s disease and experimental autoimmune encephalomyelitis, a mouse model of multiple sclerosis (Jordao et al., 2019; Mrdjen et al., 2018). In the brain, BAMs mainly localize to the meninges and CP, but not the brain parenchyma (Jordao et al., 2019; Mrdjen et al., 2018; Utz et al., 2020; Van Hove et al., 2019). Therefore, it is reasonable to speculate that BAMs might be easily exposed to pathogens or molecules from dams by being in close proximity to the vasculature (Jordao et al., 2019; Mrdjen et al., 2018; Utz et al., 2020; Van Hove et al., 2019). In contrast, the mechanism of how BAMs interact with NPCs and influence their differentiation, survival, and migration requires careful consideration. One possibility is that molecules, such as pro-inflammatory mediators, produced by BAMs are released toward ventricles and the cerebrospinal fluid (CSF), thus impacting NPCs undergoing active proliferation and differentiation in ventricular zones exposed to CSF (Supplemental Figure 7B). Further investigation is required for the mechanistic understanding of the potential interaction between BAMs and NPCs.

In this study, we provide evidence that fetal innate immunity substantially contributes to specific behavioral phenotypes induced by MIA. If both maternal and fetal immunity, independently or cooperatively, regulate diverse processes like the differentiation, migration, proliferation, and survival of NPCs, it may explain the complexity of behavior abnormalities induced by MIA. Moreover, recognizing how fetal innate immune cells contribute to the spectrum of neurodevelopmental trajectories may help us to understand the complex regulation of brain development and behaviors.

## Acknowledgements

We thank Greg Barton and his lab for the *Trif*^−/−^ mice. We thank the Saijo lab for helpful discussion. We are grateful to Ellen Robey for critical reading of the manuscript. We thank Hector Nolla and Alma Valeros for FACS sorting; Justin Choi for library preparation; and the FGL core facility for running the sequencing. We also thank Amy Sullivan (Obrizus Communications) for editing the manuscript. E. K. N. is supported by a UC Berkeley Chancellors fellowship and a UC Dissertation-Year Fellowship. K. M. G. and M. L. A. are supported by a GRFP provided by the NSF. M. T. D. is supported by a training grant (T32NS095939) and an NDSEG fellowship provided by the DoD. We thank the members of the UC Berkeley single-cell RNA-seq user group, particularly Nicholas Everetts, for their specific advice, discussion, and feedback on our analysis. This work is also supported by the Searle Scholar, the Pew Charitable Trust, and the NIH (R01HD092093) for K.S. and the Chan Zuckerberg Biohub for L.C.

## Author contribution

E. K. N. and H-C. C. equally contributed to this study and are listed in alphabetical order by their first name. K.S. conceived and designed the study. E. K. N. and H-C. C. designed and performed experiments and analyzed data. M. T. D. and M. L. A. performed experiments. P.L. and R.M. assisted with experiments. K. M. G. and L. C. advised and performed the viral experiments. E. K. N. and M. T. D. performed all the bioinformatics analysis, except for the Velocyto analysis performed by R. Z. L. K. S. wrote the manuscript with input from E. K. N., M.T.D. and H-C. C. as well as the other co-authors.

## STAR Methods

### EXPERIMENTAL MODELS AND SUBJECT DETAILS

#### Mice

All mice were maintained in specific pathogen free conditions at the animal facility at the Li Ka Shing building, University of California, Berkeley. A 12-h light/dark cycle was maintained in the housing and test rooms. Food and water were accessed *ad libitum*. Wild-type C57BL/6J (WT) mice were originally obtained from the Jackson Laboratory bred in our facilities, and *Trif*^*−/−*^ mice were originally provided by Gregory Barton. For behavioral assays, both male and female mice were used at 10-weeks old. All procedures were approved by ACUC, UC Berkeley.

### METHODS DETAILS

#### Maternal Immune Activation (MIA) by PolyI:C, LPS, and ODN

On embryonic day (E) 12.5, pregnant mice were given intraperitoneally with a single dose of either of Poly(I:C) (2 mg/kg), LPS (100 µg/kg), ODN (100 µg/kg), or saline (vehicle). Their offspring at 10-16 weeks of age were used for the behavioral assays.

#### MHV-68 and MCMV infection

Pregnant WT and *Trif−/−* mice were infected intraperitoneally at E12.5 with either MHV-68 (5 × 10^5^ plaque forming units (PFU)), MCMV (2 × 10^6^ PFU), or saline alone (vehicle). Their offspring at 2-4 months of age were used for the behavioral assays.

#### Detection of MHV-68 DNA by nested PCR

WT female mice were bred and infected intraperitoneally at E12.5 with either 200 μL 5 × 10^5^ PFU or saline. Five days post-infection (dpi), pregnant mice were sacrificed, and fetal organs (lung, spleen, and brain) were harvested. The DNA of organs was prepared by the quick-DNA miniprep kit, and analyzed for the presence of MHV-68 DNA by nested PCR targeting the gp150 gene of MHV-68. The sequences of primers were listed in the reagents and resource section.

#### Behavior Assays

Both male and female mice were used for behavior assays. The results of assays were shown either male or female offspring as indicated in the figure legends.

##### Reciprocal social interaction

A test mouse was first housed individually for at least 7 days. During the test session, an unfamiliar mouse was put into the home cage of the isolated test mouse, where it was allowed to freely interact with the test mouse. The social interactions of the test mice were recorded using a digital camera for 3 min. The time that the test mouse spent interacting with the unfamiliar mouse including sniffing, mounting, and grooming, were subsequently measured. The number of test mice observed for the aggressive behaviors including chasing, tail rattling, and biting was also recorded.

##### Three-chamber social behavior test

A three-chambered rectangular box with two dividing walls that marked the left, center, and right chambers (20 × 40 × 22 cm) was used. This box had openings that allowed a test mouse to freely move into each chamber. During the habituation phase, the mice were placed in the center chamber and allowed to freely explore for 10 min. In the sociability test, a cylindrical grid cage (7 cm in diameter and 15 cm in height) housing an unfamiliar mouse (S1), and an identical empty cage (E) were placed on opposite side chambers. The test mouse was allowed to freely explore the chambers with the S1 and E cages for 10 min to assess sociability. The movements of the test mice were recorded with a digital camera, and the time spent interacting or sniffing each cylindrical cage was measured using the Smart Video Tracking System (Panlab).

##### Marble burying assay

The fresh unscented bedding material was evenly distributed into a flat surface across the clean cage (27 × 16.5 × 12.5 cm) with a depth of approximately 5 cm. During the habituation phase, the mouse was individually placed in each cage without any marbles. After freely exploring the cage for 20 min, the mouse was temporarily removed. Then, twenty glass marbles (1.5 mm in diameter) were gently spaced in a 4 × 5 arrangement on the surface of the bedding. During the testing phase each mouse was placed in the cage with marbles and allowed to explore it for 20 min. At the end of the test, mice were removed from the cage and the number of marbles buried (>50% marble covered by bedding material) was counted.

##### Open field test

Mice were individually placed in the center of a transparent plastic box (22 × 42.5 × 21 cm) for 1 hour and allowed to freely explore the arena in the dark. Animal movement including the total travelled distance was monitored by computerized photobeam using the MotorMonitor SmartFrame System (Kinder Scientific).

##### Elevated plus maze

For the elevated plus maze test, each arm of the maze was 30 × 5 cm, and they extended out from a central platform (area 5 × 5 cm). Animals were individually placed onto the central platform and allowed to freely explore the maze for 10 min. A Smart Video Tracking System (Panlab) was used to determine the time spent in the open arms, the closed arms, and the central area during the test.

##### T-maze

The T-maze apparatus consists of a start arm perpendicular to two opposed choice arms (15 × 5 × 12 cm). Before starting the T-maze task and then throughout the entire experiment, the subject mice were food-restricted (1.5-2 g per day) to reduce their body weight to 85–90% of the free-feeding body weight. On the day of training, mice were enclosed in the stem of the start arm, habituated to the T-maze for 5 min when the door was opened, and then given two forced-correct and two forced-error trials to teach them to choose the designed goal arm. Mice then received ten training trials per day. For each trial, mice were given the choice between two arms, one of the arms being the goal arm where the sweetened-milk reward was located. If the mouse made the correct choice, it was allowed to consume the reward. The number of days taken to achieve an 80% correct response on three consecutive days was recorded to indicate the spatial learning and memory.

#### Single cell RNA-seq (scRNA-seq)

##### Tissue dissection and cell isolation

One or two E13.5 dams were sacrificed per replicate (24 hours after MIA), and individual embryos collected in ice-cold 1xPBS or DMEM F12 media (with 10% FBS, 5% HS, and 1% PS antibiotics). Embryonic tails were taken for SRY genotyping, and entire fetal brains were dissected, minced by feather blade, and triturated with a flame-polished Pasteur pipette until pieces were <1 mm in size. Whenever possible, samples were kept on ice and at 4°C during centrifugation and sorting. At least seven embryos were used per replicate; whenever possible, an equal number of male and female brain samples were pooled together. After pooling, samples were spun at 1000*xg* for 5 minutes and re-suspended in pre-warmed digestion buffer (10mL 1xHBSS with Ca^2+^ and Mg^2+^ containing Liberase and 400 U of DNAse-I). Samples were rotated end-over-end at 37°C for 10-15 minutes, with further trituration by flame-polished Pasteur pipette every five minutes. 200-400 U of DNAse-I was added. Next, the sample was strained on ice through a 100um cell strainer and washed with 4 mL of ice-cold Wash Buffer 1 (1xHBSS without Ca^2+^ and Mg^2+^ containing 200 U of DNAseI and 10% FBS) into a 50 mL conical tube. This process was repeated once more with a 70 ſm cell strainer and a fresh 50 mL conical tube. 22 mL of ice-cold Wash Buffer 2 (1xHBSS without Ca^2+^ and Mg^2+^ containing 10% FBS) was spun down at 1200*xg* for 10 minutes; cell staining followed after.

##### Myeloid cell staining and flow cytometry sorting

After centrifugation, the cell pellet was re-suspended in blocking solution (Wash Buffer 2 Fc Block at a 1:16 dilution factor) and incubated on ice for at least 10 minutes in a 2 mL lo-bind Eppendorf tube. After, Cd11b-PE-Cy7 and CD45.2-PE antibodies were added at a 1:100 dilution factor, and the sample was incubated for an additional 15 minutes on ice. 500 ſL of Wash Buffer 2 was added directly to the tub for a half-rinse; this was spun down at 800*xg* for 5 minutes and re-suspended in about 750 ſL of media (DMEM F12 without phenol red, 10% FBS, and 5% HS) per fetal brain pooled. The sample was filtered through a 40 µm cell strainer prior to sorting. For dead cell exclusion, 0.001% v/v of DAPI was added for every 100 µL of solution. An estimated 3-10×10^6^< cells per litter were stained, and on average 56,000 live- and singlet-gated Cd45^+^Cd11b^+^ was sorted into media using a BD FACSAria Fusion flow cytometer available at the Cancer Research Laboratory Flow Cytometry Facility at UC Berkeley. Cd45^+^Cd11b^+^ cells were about 1-2% of the total live cell fraction, and were not partitioned into Cd11b^hi^ or Cd11b^lo^ groups.

##### SRY genotyping

Genomic DNA from embryonic tails was isolated by digestion in 100 ſL of 50 mM NaOH until no longer visible; 25 ſL of 1 mM Tris-HCl was then added. 1 ſL of this gDNA prep was used in a fast SRY genotyping protocol using the KAPA2G Fast HotStart genotyping kit and SRY forward and reverse primers.

##### Library prep, sequencing, and raw data processing

Immediately after sorting, cells were spun down 800*xg* for 5-10 minutes, and concentrated in media to an estimated 500-1100 cells/ſL. The 10X Chromium Single Cell 3’ Reagent Kit (v3 chemistry) platform was at the Functional Genomics Lab, UC Berkeley. To maximally resolve any cellular heterogeneity in our system, we aimed for a cell capture of 9-10,000 cells per batch. The 10X Chromium Single Cell 3’ Reagent Kit (v3 chemistry) was used. All four libraries (per genotype) were pooled and sequenced on one S4 lane of the Illumina NovaSeq 6000 platform at the Vincent J. Coates Genomics Sequencing Laboratory at UC Berkeley and the UCSF Center for Advanced Technology. Sequencing reads were processed using the 10X Cell Ranger software in cluster mode.

##### Data import, filtering, and variable feature selection

The raw filtered gene expression matrices generated by the 10X Cell Ranger software was analyzed primarily using the Seurat R package (version 2.1.4) within RStudio (version 1.1.463). Genes that were present in fewer than 3 cells and cells expressing fewer than 500 genes were not used for further analysis. percent mitochondrial genes (percent.mt), and percent ribosomal genes (percent.ribo) were calculated for each cell, and only cells containing less than or equal to 15% mitochondrial genes per cell were included for downstream analysis. The data was then log normalized using Seurat’s NormalizeData function, in which the feature counts in each cell are divided by the total counts for that cell, then multiplied by a scale factor of 10,000. These values are then natural-log transformed using log1p. Next, the 10,000 most highly variable features amongst cells of each batch were selected using Seurat’s FindVariableFeatures function with the selection method parameter set to “vst”. These variable features were filtered further to the 5000 most highly variable features amongst all batches using Seurat’s SelectIntegrationFeatures. Data from each batch was merged and batch-corrected using these 5000 features with Seurat’s FindIntegrationAnchors and IntegrateData functions, using 40 dimensions and normalized using the log normalization method. These 5000 variable features were used as input for principle component analysis after additional quality control.

##### Additional cell quality control, scaling, and regression

Any low-quality cells, such as doublets remaining after filtering, were identified using the DoubletFinder tool (version 2.0.3). We then used Seurat’s CellCycleScoring function to infer the cell cycle phase of every cell; each cell received a G2M and S score (S.Score and G2M.Score, respectively) based on phase markers. The 5000 integrated features were then scaled using Seurat’s ScaleData function. Here, expression of each gene across cells was regressed against the percent of mitochondrial and ribosomal genes expressed in each cell as well as cell cycle score of each cell. The previously selected 5000 variable features were scaled on the resulting residuals These scaled residuals were used in downstream principle components analysis (PCA). We chose to keep regression of cell cycle in the analysis even though it only partially improved the effect of cell cycle on clustering.

##### Cell sex separation

The sex of every cell was identified by first merging the batches via the merge function in Seurat. The merged object for each genotype underwent sex gene expression characterization in order to infer the sex of every cell. Briefly, the lists of all genes from the Y and X mouse chromosomes were extracted from Ensembl’s biomaRt using the “ENSEMBL_MART_MOUSE” mart and the “mmusculus_gene_ensembl” dataset. The merged object data was then normalized and scaled only on features found only on the Y- and X-chromosomes. These sex gene lists were used to find all Y and X chromosome genes expressed in the dataset; only 8 total Y-chromosome features were found (*Eif2s3y, Kdm5d, Gm29650, Uty, Ddx3y, Gm29554, Gm47283*, and *Zfy1*). Meanwhile, 610 total X-chromosome genes were found; we chose *Xist* as the representative X-chromosome gene. Density histograms of gene expression were constructed for all 8 Y-chromosome genes and *Xist*; cutoff values were determined at the first minimum. For any cell that has expression of any Y-chromosome gene above the cutoff, and below the cutoff for *Xist*, were counted as male. Conversely, any cells with *Xist* expression above its cutoff, and below that of all Y-chromosome genes, were counted as female. Any cells that did not fall in this category were deemed indeterminate and removed from the study; all remaining cells had its sex assigned to the metadata.

##### Principle components selection, dataset integration, and clustering

In order to infer how many useful principal components (PCs) to include, we used Seurat’s ElbowPlot function and decided to use 25 PCs in the downstream analysis. These PCs were used to reduce the dimensionality of the dataset with Seurat’s RunPCA function. RunUMAP, FindNeighbors, and FindClusters functions were used next to visualize results. We used a k parameter of 35 for FindNeighbors, and a resolution parameter of 0.4 for FindClusters. The minimum distance parameter of RunUMAP was set to 0.4. A total of seventeen clusters were identified. The top marker genes (positive and negative) defining the clusters were found using Seurat’s FindAllMarkers function (with parameters min.pct and thresh.use at 0.25). These are considered to be differentially expressed genes. Markers for each cluster with the highest absolute value of log fold change and lowest p value were visualized with a heatmap and dotplot using Seurat’s DoHeatmap and DotPlot functions. Seurat’s DimPlot, FeaturePlot, and DotPlot functions were used to generate figures. KableExtra, ggplot2, cowplot, were also used to generate figures.

##### Gene ontology analysis

Networks of biological pathways for cluster 10 were identified with the MtGO single cell package. Genes used in the analysis consisted of the top marker genes for cluster 10 vs all other clusters as identified with Seurat’s FindMarkers function. Only genes that were expressed in at least 50% of cells in cluster 10 were included. Mitochondrial and ribosomal genes were also filtered out. Pearson’s correlation was used with MtGOsc’s ‘write.coexpresssionMatrix’ function to generate a matrix of coexpression between each pair of marker genes for all cells in cluster 10. Pairwise correlations of genes with the top 5% of Pearson’s r values were selected as network edges. Edges weighted by their r values were used to generate a network of biological pathways as previously described (Yuan et al., 2016). For this, a gene ontology (GO) dictionary downloaded from the Mouse Genome Informatics database provided all possible GO terms for each gene. Genes and their GO terms were then clustered into functional modules with overlapping GO terms using the MTGO algorithm. A minimum of 2 genes was allowed to create a cluster defining a biological pathway and the max gene ontology parameter was set to 125. VisNetwork and iGraph were used to generate network plots.

##### RNA velocity analysis

RNA velocity analysis was done using the Velocyto package. Fastq files were trimmed and mapped using STAR to the mouse reference genome (*mm10*). From the annotation files, intronic and exonic information was extracted. From this information, ratios of the spliced and unspliced variants of each gene in every cell is known; changes in these ratios for a gene in a cell allow for inferences in cell state changes among the population. Arrows point to the position of the future state. The data presented here comes from the WT (and not *Trif*^*−/−*^) portion of the dataset. The arrow embeddings were plotted and visualized by tSNE.

#### Immunofluorescence

The 30 μm-thick sagittal brain sections were blocked by incubating in PBST (0.1% Triton X-100 in PBS) containing 5% normal donkey serum and then incubated with primary anti-CD206 (1:100), anti-P2Y12 (1:300), and anti-S100a8/9 (1:100) antibodies overnight at 4°C. After washing, the brain sections were incubated with the corresponding secondary antibodies conjugated with Alexa Fluor® 488, Alexa Fluor® 594, and Alexa Fluor® 647 at room temperature for 2 hr, stained with iamidino-2-phenylindole (DAPI, 1:10000) to label nuclei, and mounted in fluoromount-G Mounting Medium. Images were acquired using a confocal microscope (LSM 700; Carl Zeiss). The numbers of CD206-positive, P2Y12-positive, and CD206- and S100a8/9-double positive cells in the choroid plexus, meninges, and the fetal cortex were measured with Fiji. The integrated density of intracellular S100a8/9 immunoreactivity in the CD206 positive cells in the choroid plexus was measured. For each mouse, at least 5-7 images of the desired brain regions were sampled and averaged.

