## Supplementary material for "Fetal innate immunity contributes to the induction of atypical behaviors in a mouse model of maternal immune activation": Supplemetal Figures

**Supplemental information**

**Supplemental Figure Legends**

**Supplemental Figure 1. MIA-induced behavior abnormalities. A.** A simplified diagram of TLR signaling is shown. TLR3 is expressed on the endosomal membrane and signals through TRIF to activate the transcription of anti-viral response genes. TLR9 is also expressed on the endosomal membrane and signals through MyD88 to activate the transcription of pro-inflammatory mediators. TLR4 is expressed on the plasma membrane and signals through both TRIF and MyD88. **B.** A table that summarizes the behavior assays used in this study is shown. **C.** Hyperactivity-like behavior was determined by the open field assay and is shown as total distance (in, inches) that the mice moved. Male offspring from control- (Veh, black circles, N=13), Poly I:C- (red squares, N=15), and LPS- (blue triangles, N=9) MIA-treated dams were tested. **D.** The degree of anxiety-like behavior was determined by the elevated plus maze assay and is shown as the time spent in closed arms (sec, seconds). Male offspring from vehicle control- (Veh, black circles, N=4), Poly I:C- (red squares, N=10), and LPS- (blue triangles, N=7) MIA-treated dams were tested. **E.** Memory and learning was tested by T-maze and is shown as the average days to acquisition. Offspring from control- (Veh, black circles, N=5), Poly I:C- (red squares, N=13), and LPS- (blue triangles, N=7) MIA-treated dams were tested. **F.** Social interaction time (Δsec) with a stranger mouse determined by the three-chamber social interaction is shown. Offspring from vehicle control- (Veh, open circles, N=7) and CpG ODN- (green diamonds, N=11) MIA-treated dams were used. **G.** The degree of repetitive restricted behavior was determined by the number of marbles buried in the marble burying assay. Offspring from vehicle control- (Veh, open circles, N=7) and CpG ODN- (green diamonds, N=13) MIA-treated dams were used. **H.** Memory and learning was tested using the T-maze assay and is shown as the average days to acquisition. Offspring from vehicle control- (Veh, open circles, N=7) and CpG ODN- (green diamonds, N=12) MIA-treated dams were used. **I.** The degree of anxiety-like behavior was determined by the elevated plus maze and is shown as the time spent in closed arms (sec). Offspring from vehicle control (Veh, black circles, N=7) and CpG ODN- (green diamonds, N=13) MIA-treated dams were used. **J.** Hyperactivity-like behavior was determined by the open field assay and is shown as the total distance (in, inches) that the mice moved. Offspring from vehicle control (Veh, open circles, N=7) and CpG ODN- (green diamond, N=12) MIA-treated dams were used. **B-I.** Data indicate individual animals and averages. Error bars show S.E.M. **K.** The number of offspring from the indicated MIA models exhibiting aggressive behavior (see methods for details) during the reciprocal social interaction assay is shown. **L.** A summary of the behavior abnormalities observed in offspring from the CpG ODN-MIA model compared to vehicle control is shown. **C-K.** Male offspring from control-MIA (Veh, N=5), Poly I:C-MIA (N=12), LPS-MIA (N=8), and CpG ODN -MIA (N=12) and its control (Veh, N=7) were used. * indicates $p<0.05$, ** indicates $p<0.01$, and *** indicates $p<0.001$.

**Supplemental Figure 2. Requirement of maternal and fetal TRIF signaling in other behaviors. A and D.** Memory and learning was tested using the fear conditioning assay and is shown as the percentage of freezing. **B and E.** The degree of anxiety-like behavior was determined by the elevated plus maze assay and is shown as the time spent in the closed arms (sec, seconds). **C and F.** Hyperactivity was determined using the open-field assay and is shown as the total distance (in, inches) that the mice moved. **A-C.** Offspring with a genotype of *Trif*$^{+/-}$ (Figure 2 A) from control- (Veh, black circles, N=11) and Poly I:C- (red squares, N=8) MIA-treated dams were used for these assays. **D-F.** Offspring with a genotype of *Trif*$^{-/-}$ (see Figure 2D) from control- (Veh, black circles, N=5) and Poly I:C- (red squares, N=7) MIA-treated dams were used. Data indicate individual animals and averages. Error bars show S.E.M. Asterisk (*) indicates $p<0.05$, and n.s. indicates not significant.

**Supplemental Figure 3. Characterization of the batch effects and *Fcrls*$^+$ clusters. A.** A table summarizing the data from a total of eight batches that were used for our analyses. **B.** The number of unique molecular identifiers (nUMIs) per cell in each batch post quality control is shown on a violin plot. UMIs correspond to barcoded mRNA captured per cell and the identification numbers of the batches are shown in A. **C.** The expression levels of *Fcrls* in the clusters are shown on the histogram. Cluster IDs are indicated on the y-axis and the degree of expression is indicated on the x-axis. The expression patterns of BAM- (**D**) and microglia- (**E**) specific genes (indicated at the top) in the cell clusters are shown as violin plots.

**Supplemental Figure 4. Characterizations of clusters and contributions of male and female cells. A.** The contribution of cells from the offspring of Poly I:C- (red) and vehicle- (blue) MIA-treated mice to each cluster is shown. The % of total cells per treatment is on the y-axis and Cluster IDs are indicated on the x-axis. **B.** The contribution of the individual batches indicated in Supplemental Figure 3A to each of the clusters are shown. **C**. UMI plot after downsampling. Broken circles indicate Clusters 9 and 10. **D.** Contribution of male and female cells to the UMAP plot in Figure 3B is shown. Cluster IDs are the same as in the original figure. **E.** The percentages of male and female cells in Clusters 9 (top) and 10 (bottom) are displayed.

**Supplemental Figure 5. Characterization of Clusters 9 and 10. A.** UMAP plots indicating stages of the cell cycle. Cluster IDs are shown on the left. **B**. Inter-molecule connection plot described in Figure 5B and C is shown. **C.** Volcano plot representing the differential gene expression of Cluster 9 compared to Clusters 0 (BAMs)(left) and 1(microglia) (right). Orange dots represent differentially expressed genes in Cluster 9 compared to Clusters 0 (left) and 1 (right) (upregulated gene cutoff is log2 fold change > 1.5 and $p$ $adj$ < 0.01; downregulated gene cutoff is log2 fold change < -1.5 and $p$ $adj$ < 0.05).

**Supplemental Figure 6. Quantification of BAMs and microglia. A.** Representative images of anti-CD206 antibody staining of meninges from the fetal brains of E13.5 offspring from vehicle-treated (left) and Poly(I:C)-treated (right) dams are shown. Scale bars are 50 μm. **B**. Quantification of the images shown in A is shown. Data is shown as average cell numbers from 5 to 6 images of serial sections from control (Veh, black circle, N=4) and Poly(I:C)-MIA induced (red rectangle, N=3) of meninges. **C.** Representative images of anti-P2Y12 antibody staining of fetal brains of E13.5 offspring from dams treated with vehicle- (left) and Poly(I:C)- (right) MIA. Scale bars are 50 μm. **D**. Quantification of the images shown in B is shown. Average cell numbers from 5 to 7 images of serial sections of the CP region from control (Veh, black circle, N=4) and Poly(I:C)-MIA induced (red rectangle, N=3) are shown. **E**. Representative images of anti-P2Y12 antibody staining of the cortical parenchyma of fetal brains from E13.5 offspring of dams treated with vehicle- (left) and Poly(I:C)- (right) MIA. Scale bars are 50 μm. **F**. Quantification of the images shown in F is shown. Average cell numbers from 5 to 6 images of serial sections of the cortical parenchyma from control (Veh, black circle, N=4) and Poly(I:C)-MIA induced (red rectangle, N=3) are shown. n.s. indicates not significant.

**Supplemental Figure 7. Nested PCR analysis of MHV-68 and additional behavior assays of offspring from virus-infected dams. A.** Results of nested PCR for MHV-68 viral DNA in fetal tissues following MIA are shown. **B.** The working model is shown. At the steady-state, microglia cells localize in the brain parenchyma, and BAMs localize in Choroid plexus. Upon sensing Poly(I:C) or virus, BAMs in the CP are activated and produce pro-inflammatory mediators. These molecules may disrupt normal neurogenesis and lead to abnormal behaviors.

A.

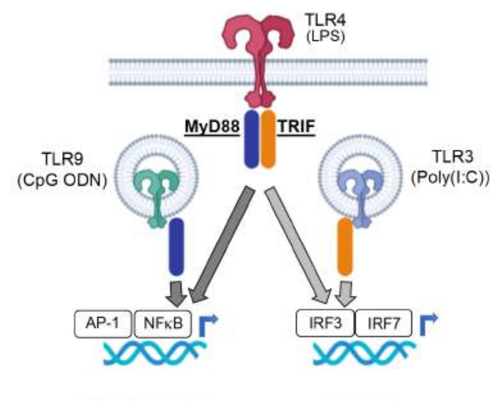


B

| Behavior | Assays |
|---|---|
| Sociability | Reciprocal social interaction assay<br>3-chamber interaction assay<br>Food preference social transmission assay |
| Repetitive restricted behavior | Marble burying assay<br>Open field assay* (excessive grooming) |
| Learning/Memory | T-maze assay<br>Fear conditioning assay |
| Anxiety | Elevated plus maze assay<br>Open field assay* (center vs peripheral ratio) |
| Hyperactivity | Open field assay* (general and total activity) |

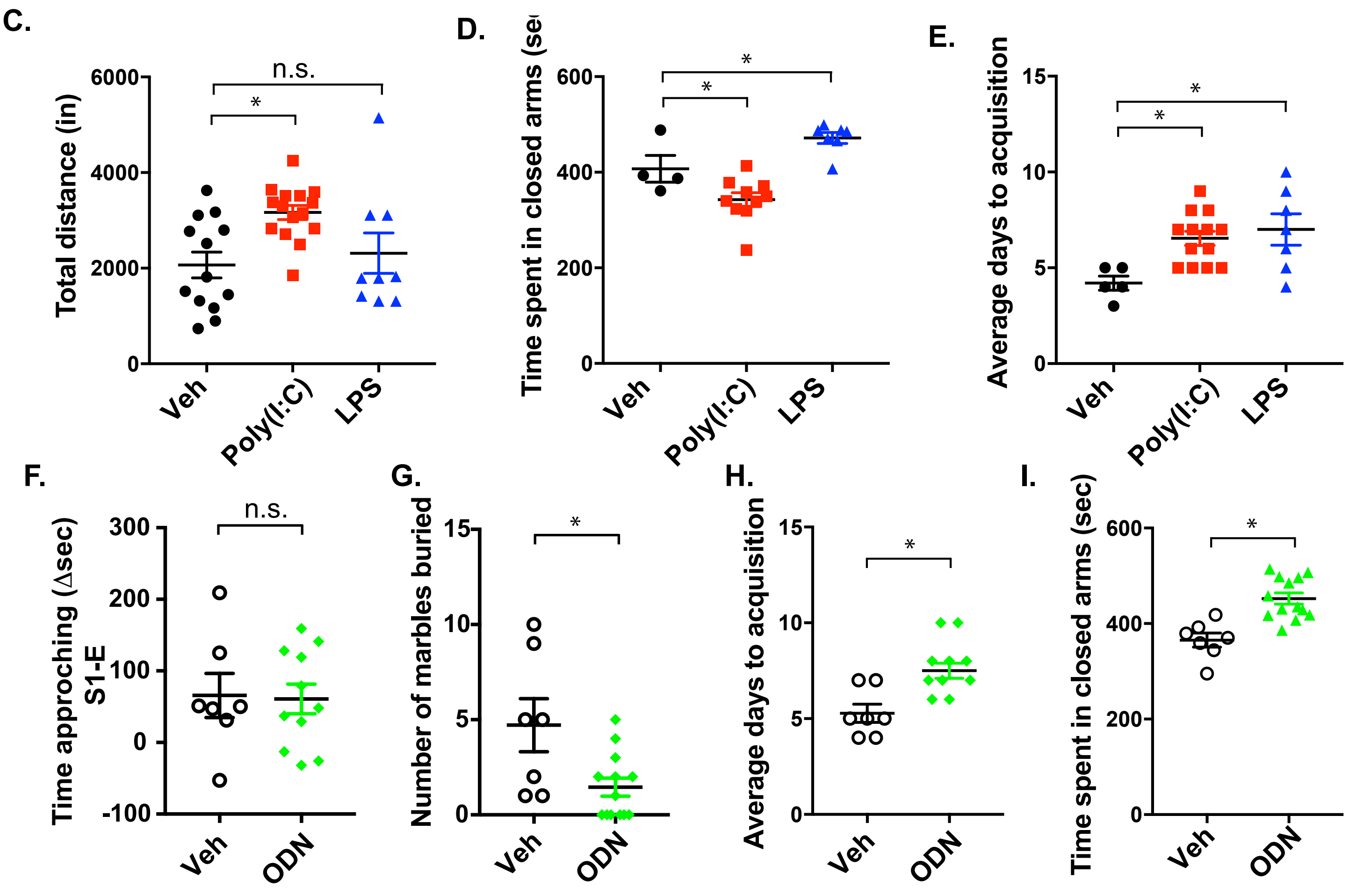


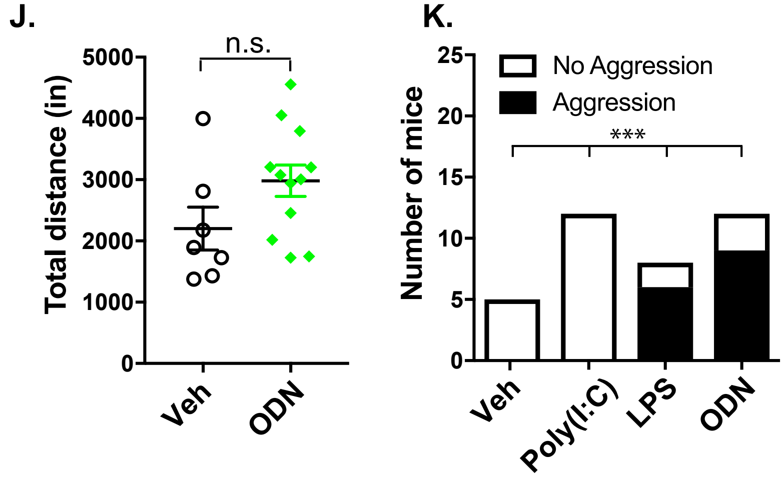


L.

| Behavior | CpG ODN |
|---|---|
| Sociability | Aggression |
| Restricted | ➡ |
| Learning/memory | ⬇ |
| Anxiety | ⬆ |
| Hyperactivity | ➡ |

Supplemental Figure 1

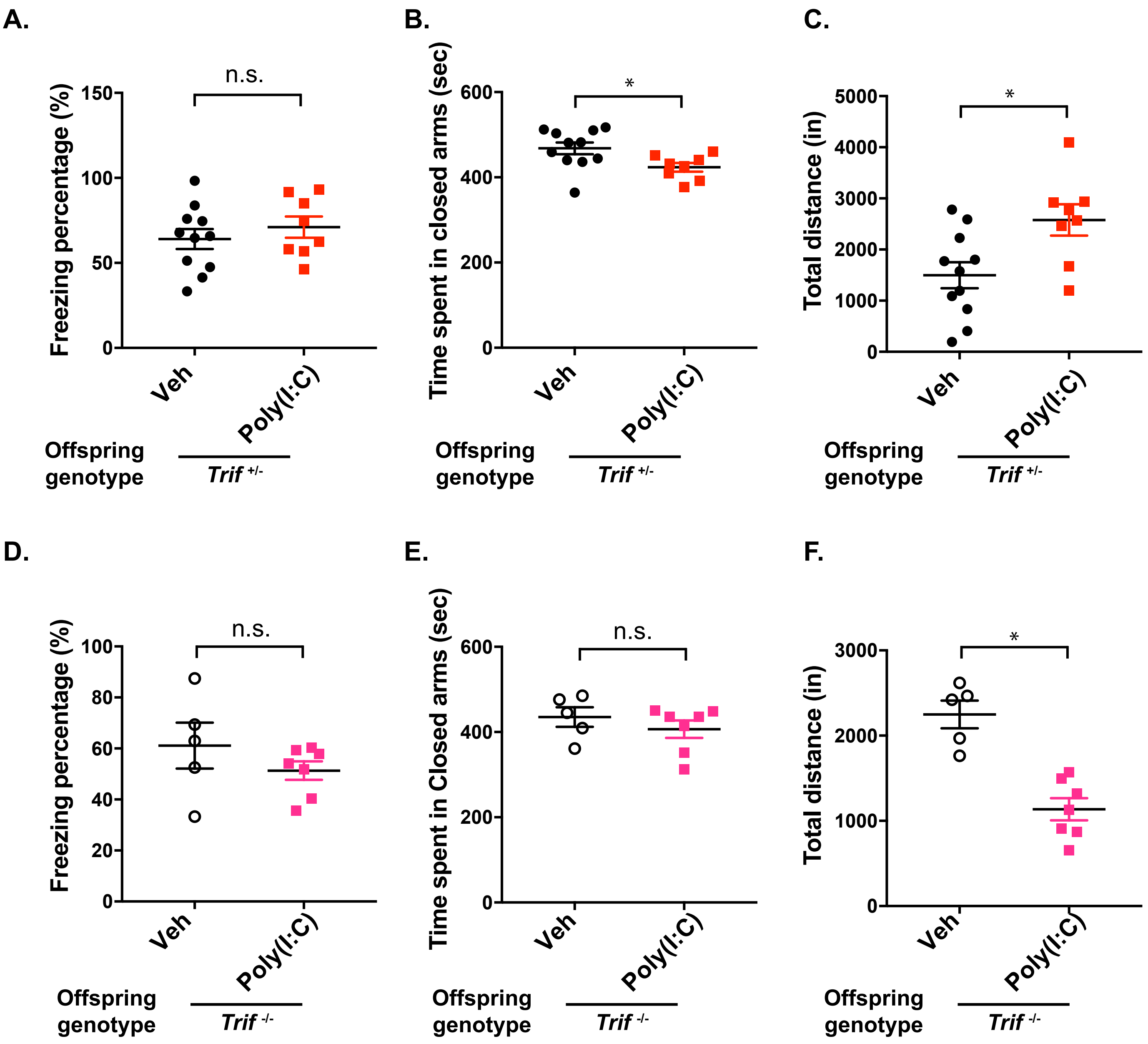


Supplemental Figure 2

**A.**

| | Batch ID | | | | | | | |
|---|---|---|---|---|---|---|---|---|
| | A | B | C | D | E | F | G | H |
| | Trif KO | | | | WT | | | |
| **Treatment_replicate** | Poly(I:C) #1 | Poly(I:C) #2 | Vehicle #1 | Vehicle #2 | Poly(I:C) #1 | Poly(I:C) #2 | Vehicle #1 | Vehicle #2 |
| **Number of litters** | 2 | 2 | 1 | 1 | 1 | 1 | 1 | 1 |
| **Total embryos** | 19 | 18 | 9 | 7 | 10 | 9 | 8 | 8 |
| **Total male embryos** | 5 | 3 | 3 | 1 | 6 | 3 | 4 | 4 |
| **Total female embryos** | 14 | 15 | 6 | 6 | 4 | 6 | 4 | 4 |
| **Total cells (post quality control)** | 4463 | 5545 | 3708 | 5832 | 4696 | 3717 | 7045 | 3600 |

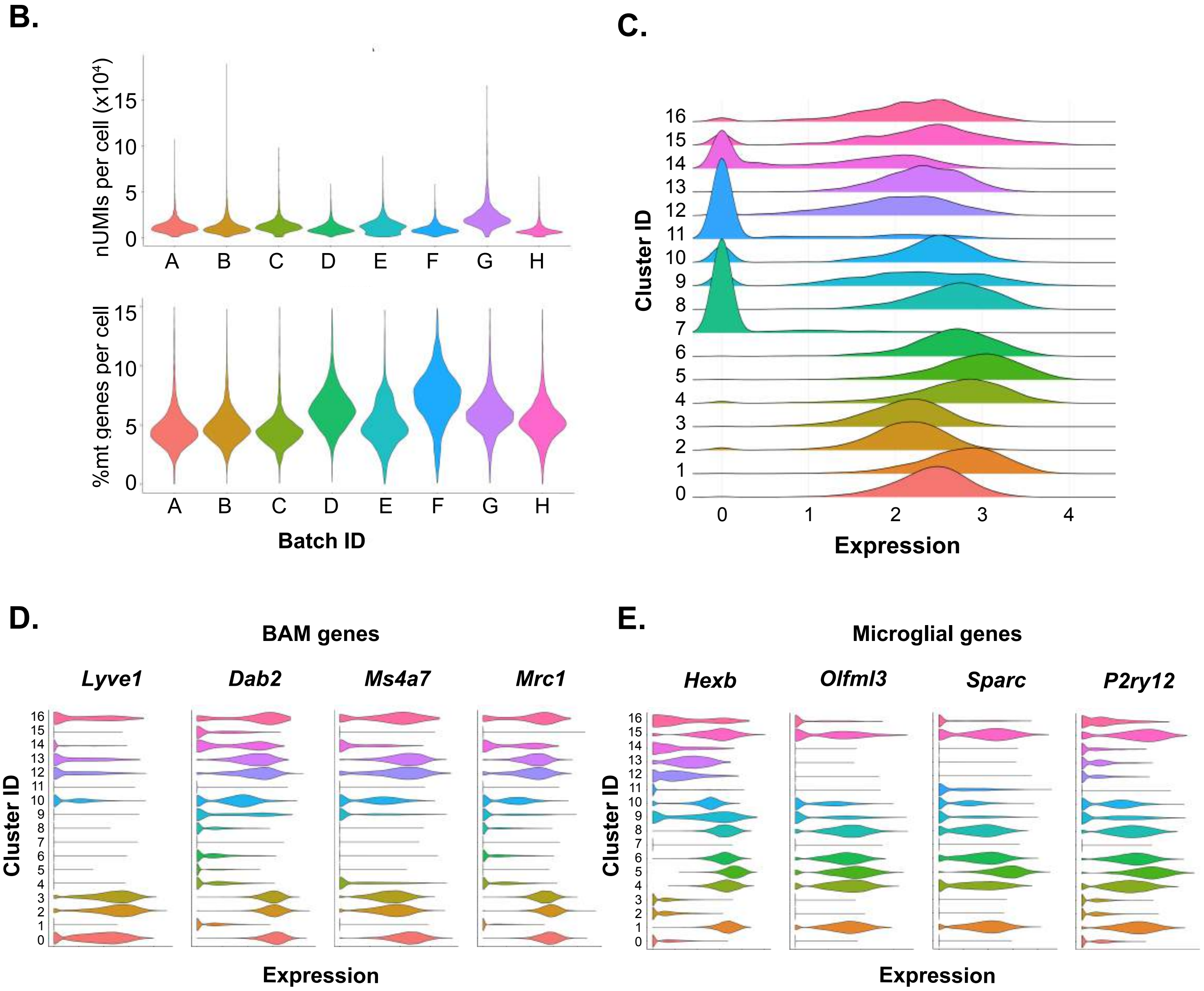


Supplemental Figure 3

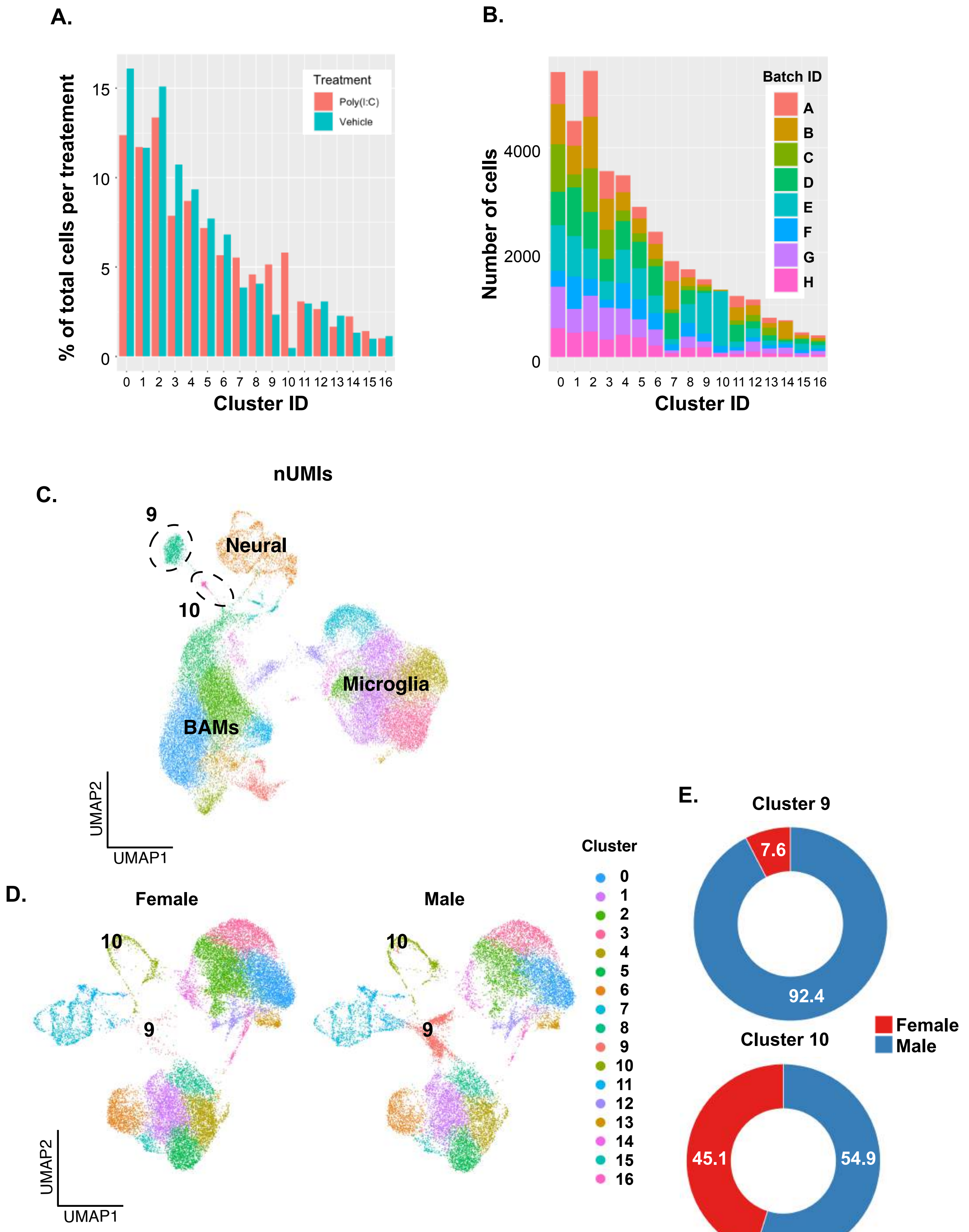


Supplemental Figure 4

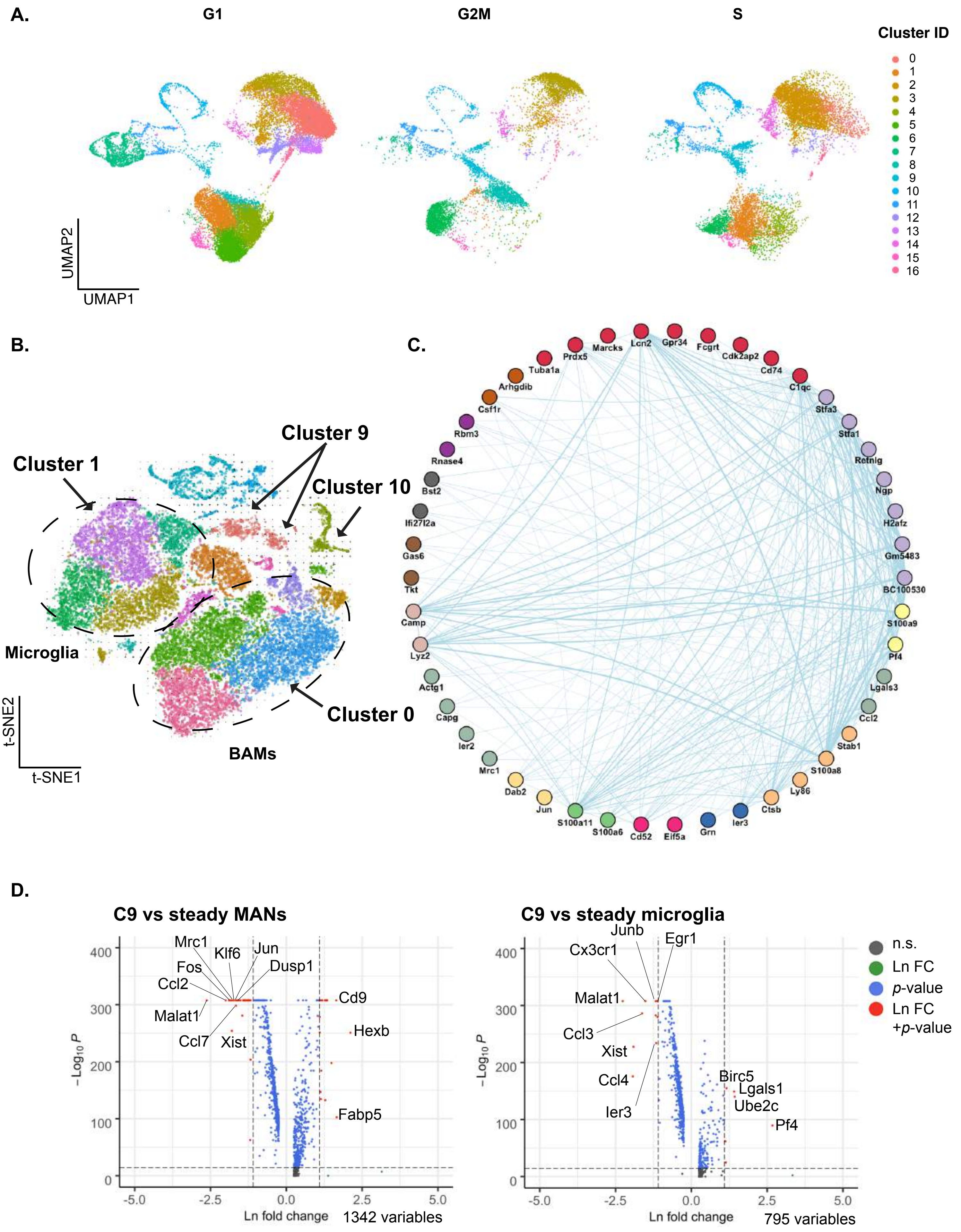


Supplementary Figure 5

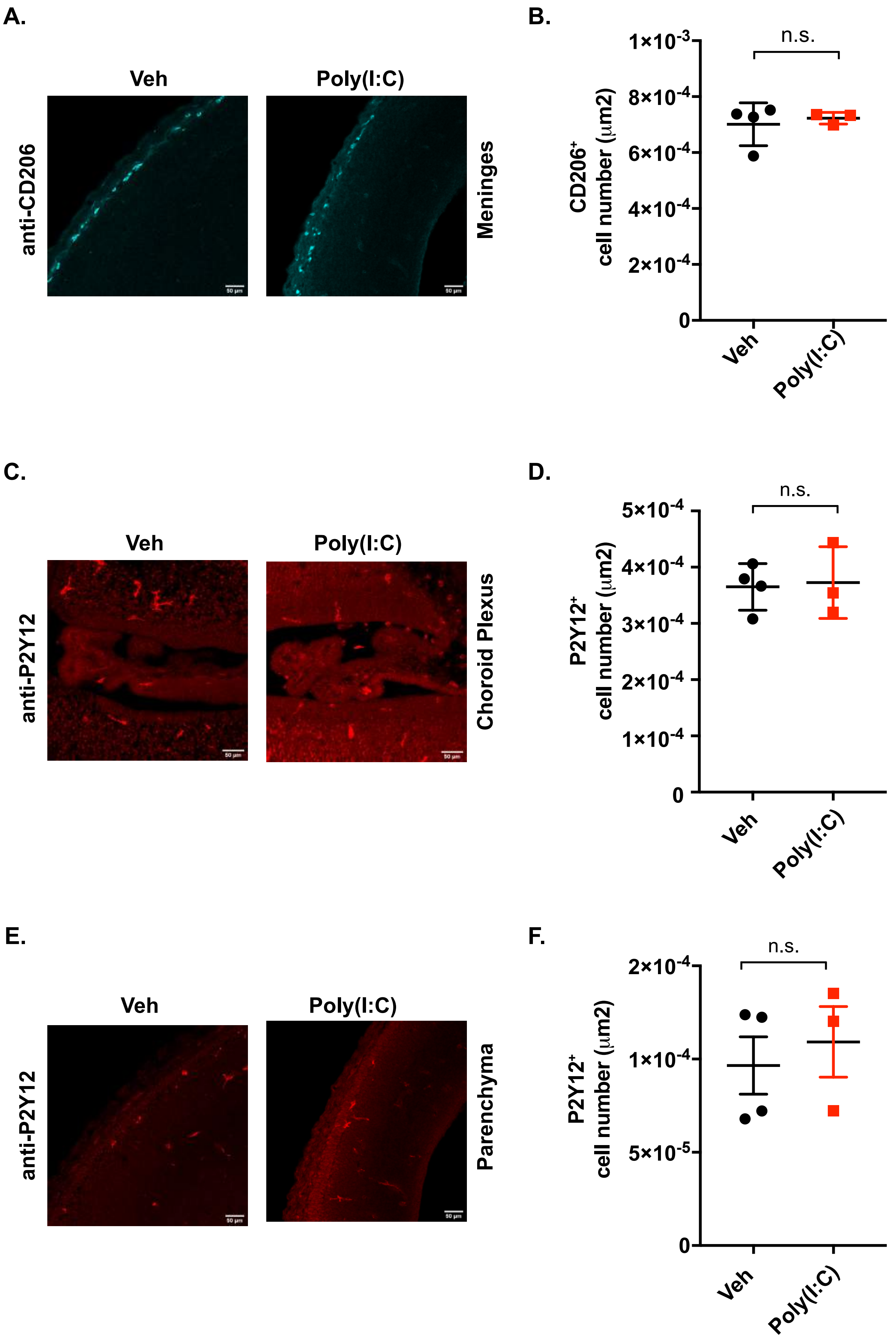


Supplemental Figure 6

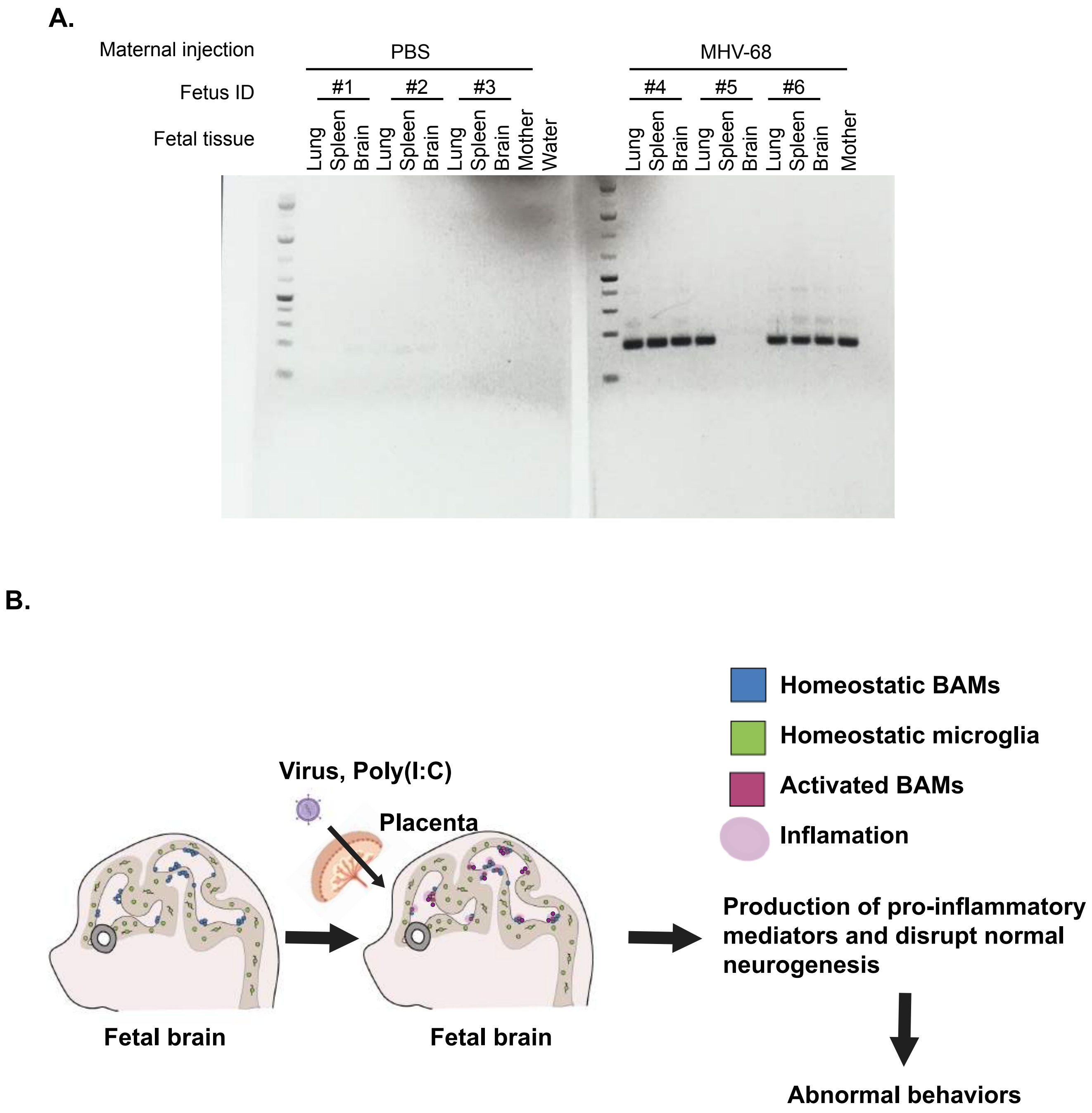
A.
Maternal injection
PBS
MHV-68
Fetus ID
#1
#2
#3
#4
#5
#6
Fetal tissue
Lung
Spleen
Brain
Mother
Water
B.
Homeostatic BAMs
Homeostatic microglia
Activated BAMs
Inflamation
Virus, Poly(I:C)
Placenta
Fetal brain
Fetal brain
Production of pro-inflammatory mediators and disrupt normal neurogenesis
Abnormal behaviors
